# Key Regulatory Elements of the TGFβ-LRRC15 Axis Predict Disease Progression and Immunotherapy Resistance Across Cancer Types

**DOI:** 10.1101/2024.11.22.624939

**Authors:** Claire M. Storey, Michael Cheng, Mohamed Altai, Julie E. Park, Julie Tran, Smiths S. Lueong, Daniel Thorek, Liqun Mao, Wahed Zedan, Constance Yuen, Alexander Ridley, Marija Trajovic-Arsic, Ken Herrmann, Sumit K. Subuhdi, Bilal A. Siddiqui, Katharina Lückerath, Jens Siveke, Robert Damoiseaux, Xia Yang, David Ulmert

## Abstract

Transforming growth factor-beta (TGFβ) has dual roles in cancer, initially suppressing tumors but later promoting metastasis and immune evasion. Efforts to inhibit TGFβ have been largely unsuccessful due to significant toxicity and indiscriminate immunosuppression. Leucine-rich repeat-containing protein 15 (LRRC15) is a TGFβ-regulated antigen expressed by mesenchymal-derived cancer cells and cancer-associated fibroblasts (CAFs). In preclinical studies, ablation of TGFβ-driven LRRC15+ CAFs increased tumor infiltration of CD8+ T cells. However, the underlying pathobiological mechanisms prompting TGFβ’s upregulation of LRRC15 expression are unclear. Using an integrated approach combining functional compound screening with single-cell RNA sequencing, we reveal key genomic features regulating TGFβ’s ability to increase LRRC15 expression on cancer cells. Construction of gene regulatory networks converged our analyses on four key genes—*MMP2, SPARC, TGF*β*R2,* and *WNT5B*—central to TGFβ-induced LRRC15 pathobiology. Validation of these genes in cell models and their use in predicting immunotherapy responses highlight their potential in refining immunotherapy strategies and personalizing co-treatment options.

## INTRODUCTION

The transforming growth factor-beta (TGFβ) signaling pathway, recognized for its dual role in cancer biology, acts as a tumor suppressor in early-stage malignancies but promotes progression, metastasis, and immune evasion in aggressive cancers and advanced stages of disease (1–3). This paradoxical behavior complicates targeting of TGFβ in oncology, as its functions vary widely from tumor suppression to the facilitation of epithelial-mesenchymal transition (EMT) and therapy resistance (4–6). TGFβ also plays a role in both primary and acquired resistance to treatment; signaling initially contributes to an immunosuppressive microenvironment that shields emerging tumors from immune surveillance. During acquired resistance, TGFβ signaling intensifies, enabling cancer cells to evade ongoing therapies by promoting invasiveness, metastasis, and maintaining an immunosuppressive microenvironment (7–9).

Leucine-rich repeat-containing protein 15 (LRRC15) has been recognized for its role as surrogate marker for TGFβ signaling and its association with immunosuppression and therapy resistance in aggressive TGFβ-mediated malignancies, notably in treatment-resistant osteosarcoma (10). Its predominant association with cancer-associated fibroblasts (CAFs) in the tumor microenvironment (TME) of immune-excluded, metastatic, and aggressive primary tumors further underscores its therapeutic relevance (11). Investigations into the TGFβ-driven TME have identified that LRRC15 is a part of an 11-gene signature that correlates with immune checkpoint therapy (ICT) resistance in patients (12–14). Furthermore, recent data from engineered mouse models demonstrated that targeted depletion of LRRC15+ CAFs markedly diminishes tumor fibroblast content and augments CD8+ T cell efficacy (13). Our previous independent verification using a novel radiotheranostic antibody-based approach confirmed that radioimmunotherapeutic targeting of LRRC15+ cells significantly decreased tumor burden and stopped disease progression, along with suppression of genes linked to TGFβ-driven immunotherapy resistance (15).

In this study, we evaluated the complex interplay of TGFβ signaling and LRRC15 expression within mesenchymal stem cell (MSC)-derived tumors. We show a bifurcated cellular response to TGFβ activity; certain tumor cells undergo a rapid induction of LRRC15, while expression is unaltered in others. To further explore the transcriptional dynamics observed in TGFβ-responsive LRRC15+ cell models, we comprehensively investigated how this differential expression of LRRC15 is indicative of TGFβ-mediated tumor pathobiology. Integration of high-throughput screening (HTS) to identify small molecules that modulate TGFβ-mediated LRRC15 induction, along with single-cell RNA sequencing (scRNAseq) to identify the gene networks and key regulators of LRRC15 inducibility, identified four genes—*TGF*β*R2*, *SPARC*, *MMP2*, and *WNT5B*—that activate TGFβ-mediated LRRC15 expression. Further evaluation in patient tumor cohorts revealed that these activating genes, alongside LRRC15, are a prognostic determinator for patient response to immunotherapy and the progression of aggressive malignancies.

## RESULTS

### TGF**β**1 induces LRRC15 expression in cancer cells

Previous studies have shown that TGFβ1 induced LRRC15 expression in MSCs under supraphysiological conditions over extended periods of time (16). In this study, we investigated the response of LRRC15 expression in cancer cells to physiologically relevant TGFβ1 concentrations, simulating conditions within the TME (17). Nine LRRC15+ cancer cell lines (CALU1, KASUMI2, SAOS2, U2OS, HUO9, NCI-H196, U118, U87, RPMI7951) (15) underwent 24-hour incubation in reduced-serum media prior to reintroduction of TGFβ1 at concentrations from 0 to 10 ng/mL (Figure 1A). Subsequently, plasma membrane-associated LRRC15 was detected via flow cytometry using the AlexaFluor-647 labeled anti-LRRC15 IgG1 antibody (DUNP19).

**FIGURE 1.**
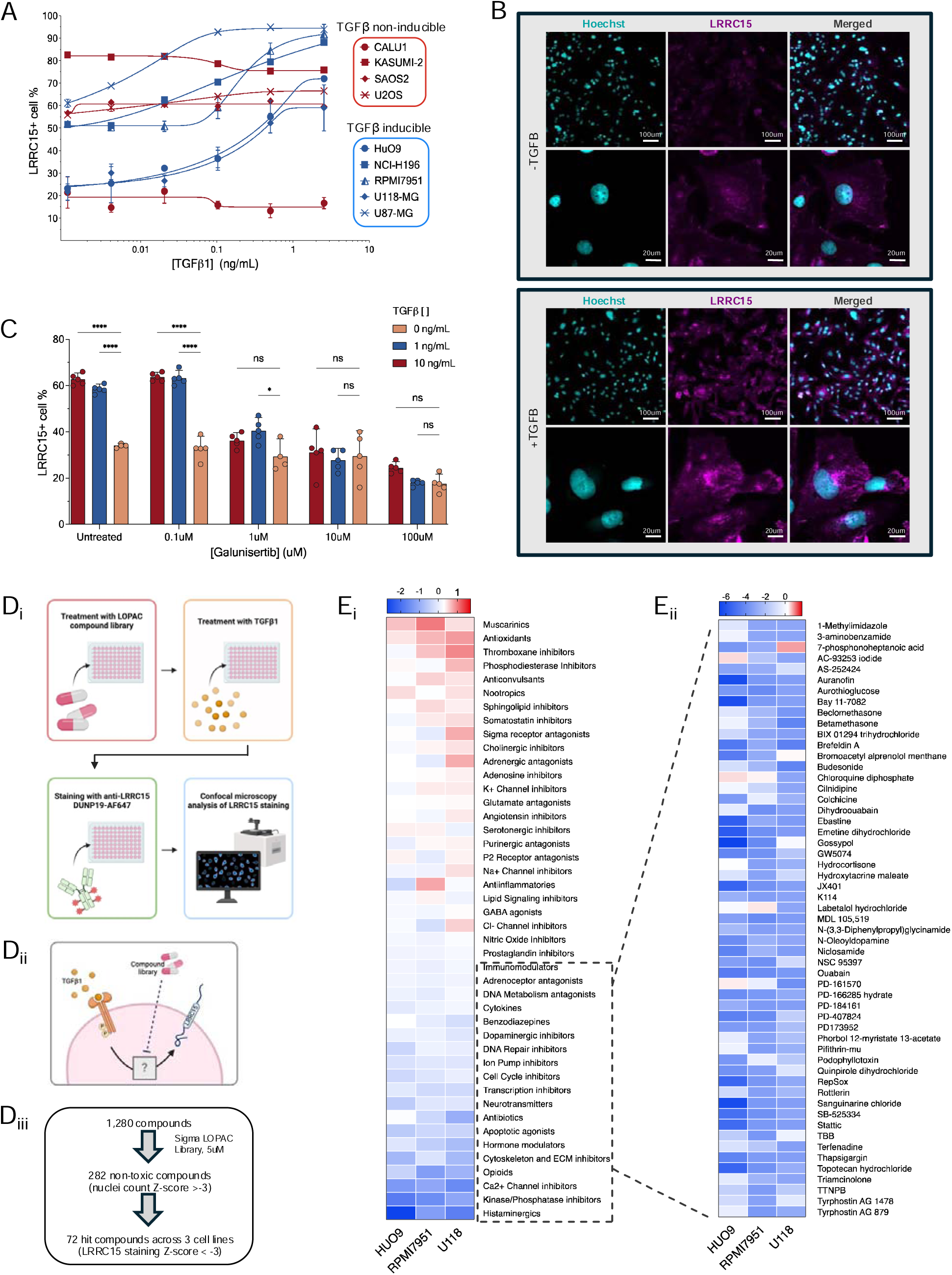
Dissection and modulation of the TGFβ-LRRC15 axis in select cancer cell lines. **(A)** LRRC15+ cancer cell lines were exposed to TGFβ1 concentrations from 0 to 10 ng/mL following 24 hours of serum starvation to simulate physiological conditions. Flow cytometry analysis (n = 3-4 per cell line) shows variable LRRC15 expression responses across cell lines, with minimal changes observed in CALU1, KASUMI-2, SAOS2, and U2OS (TGFβ non-inducible, red), and significant upregulation in HUO9, NCI-H196, RPMI7951, U118, and U87 (TGFβ inducible, blue) (pC<C0.0001). LRRC15 staining is represented as mean ± SD. **(B)** Confocal microscopy of LRRC15 expression in TGFβ-responsive cell line RPMI7951 after treatment with or without TGFβ. Cells were stained with Hoechst (nuclei, cyan) and anti-LRRC15 IgG1-AlexaFluor-647 (magenta). **(C)** Pretreatment with the TGFβ receptor inhibitor Galunisertib significantly suppressed TGFβ-induced LRRC15 expression in a dose-dependent manner (pC<C0.0001), confirming pathway specificity. Data is expressed as average LRRC15 expression (by confocal microscopy) across U118, RPMI7951, and HUO9 cells. **(D)** Schematic of high-throughput screening workflow. **(i)** LRRC15+ cells were pre-treated with the LOPAC1280 compound library, followed by TGFβ1 reintroduction and analysis of LRRC15 expression using confocal microscopy. **(ii)** Compounds were classified based on mechanism of action, and the effects on LRRC15 expression were recorded. **(iii)** 1,280 compounds were screened, and 282 non-toxic compounds were identified (cell count Z-score > –3). Of these, 72 compounds were hits, reducing LRRC15 expression below 3 standard deviations from the plate mean. **(E)** Heatmap (red = increase, blue = decrease) of LRRC15 staining Z-scores across compound classes (left heat map) and all hit compounds (right heat map) after screening with LOPAC1280 compound library.

These assessments revealed variable responses to TGFβ, with minimal changes in LRRC15 expression levels in CALU1 (percent change in LRRC15 staining = –21.94%, p = 0.2805), KASUMI2 (–8.16%, p = <0.0005), SAOS2 (+4.89%, p = 0.0517) and U2OS (+18.98%, p = 0.0008), while pronounced increases were noted in HUO9 (+210.26%, p = 0.0220), NCI-H196 (+70.82%, p = <0.0005), U118 (+173.06%, p = <0.0005), U87 (+53.65%, p = <0.0005), and RPMI7951 (+76.45%, p = 0.0038) (Figure 1A). For subsequent analyses, the latter cell lines were deemed “TGFβ-responsive” for their ability to upregulate LRRC15 after treatment with exogenous TGFβ1. Among the TGFβ-responsive cell lines, we focused on three models representing aggressively growing malignant tissues from different anatomical origins: HUO9 (osteosarcoma), RPMI7951 (malignant melanoma), and U118 (glioblastoma multiforme). Using a confocal microscopy-based approach with an AlexaFluor-647-conjugated DUNP19, we confirmed a significant increase in LRRC15 expression in cells treated with TGFβ compared to controls as shown for the RPMI7951 cell line (Figure 1B). Furthermore, pretreatment with the TGFβ receptor inhibitor galunisertib before addition of TGFβ1 effectively suppressed LRRC15 induction in HUO9, RPMI7951, and U118 cells, demonstrating the role of TGFβ in modulating LRRC15 expression (Figure 1C).

Next, we sought to annotate the molecular mechanisms impacting TGFβ’s regulation of LRRC15 (Figure 1D_i__-iii_). We conducted a high-throughput screen using the LOPAC1280 library, which comprises 1,280 well-annotated small molecules spanning various mechanisms of action and drug classes (Figure 1D_ii__-iii_). LRRC15+ cells were incubated in reduced-serum media to deplete endogenous cytokines before treatment with the compound library and TGFβ (Figure 1D_i_). Following compound incubation, cells were treated with TGFβ and assessed for LRRC15 expression via confocal microscopy (Figure 1D_i_). All screened compounds were categorized by class and mechanism, and the normalized results were presented as their average effect on LRRC15 expression (Figure 1E_i_). Compounds that significantly inhibited TGFβ-driven LRRC15 expression, defined as those reducing the integrated intensity of LRRC15 by more than three standard deviations (SD) from the plate mean, were identified as hits (Figure 1E_ii_). In addition, compounds were assessed for toxicity; those reducing cell viability >3 SD from the mean were excluded from subsequent analyses. Ultimately, we identified 26, 22, and 24 hits in the HUO9, RPMI7951, and U118 cell lines, respectively (Supplemental Table 1).

Established TGFβR2 antagonists such as SB-525334 and RepSox further validated the specificity of the assay and the mechanism by which LRRC15 is upregulated (Supplemental Table 1). Notably, various histaminergics, antivirals, and hormone modulators, for example chloroquine, Tenidap, and hydrocortisone, also reduced TGFβ-induced LRRC15 expression, underscoring the diverse roles of TGFβ in inflammation and immune modulation via diverse biological pathways (Figure 1E). Other compound classes, including potassium channel inhibitors and angiotensin inhibitors, increased LRRC15 expression above baseline (Figure 1E). These compound hits were particularly intriguing given LRRC15’s previously reported role in SARS-CoV-2 infection and structural similarity to the angiotensin-converting enzyme ACE2 (18), suggesting some utility in preventing viral infection. All gene targets from the effective compounds within our screen were compiled for further analysis.

### Single-cell RNA sequencing of TGF**β**-responsive cell lines reveals distinct LRRC15-related signatures

After identifying compounds that could disrupt the TGFβ-LRRC15 pathway, we performed scRNAseq on the cell lines utilized in the compound screen. In addition to the screened cell lines, three TGFβ non-responsive cell lines (CALU1, KASUMI2, SAOS2) were also sequenced and used in subsequent differential gene and pathway analyses (Figure 2A). To reduce gene expression sparsity, we used a high-dimensional weighted gene co-expression network analysis (hdWGCNA) metacell aggregation approach (19), which aggregates neighboring cells based on gene expression similarity and averages their expression to create metacells that are robust to scRNAseq dropouts Figure 2A. This enabled us to identify individual differentially expressed genes (DEGs) and transcriptomic pathways that were exclusive to TGFβ-responsive cells. While the lower dimension representation of the gene expression data did not separate the six cell lines by TGFβ inducibility (Figure 2B), we identified sets of DEGs with consistent effects in TGFβ-responsive cells (Figure 2C). Key cell proliferation and non-canonical TGFβ signaling genes, including TGFβ activator *LTBP1* (latent-transforming growth factor beta-binding protein 1) and Wnt ligand *WNT5B*, were upregulated in TGFβ-responsive cells. Interestingly, T-cell modulators and inflammatory driver genes such as *IL-7R* (interleukin-7 receptor) (20) were downregulated in TGFβ-inducible cell lines, providing insights into the potential immunoregulatory mechanisms within the TGFβ-LRRC15 pathway (Figure 2C). Pathway enrichment analysis of upregulated genes in the TGFβ-responsive cells showed enrichment of apoptotic processes and cytokine signaling within immune pathways, while downregulated DEGs were associated with translation and ribosomal function (Figure 2D). Furthermore, the DEG effect, calculated using metacell expression, followed a similar pattern in single cell expression UMAP plots. Several genes such as *SPARC*, *WNT5B*, *MMP2*, and *EID1*, showed a stark upregulation of expression within TGFβ-responsive cells (Figure 2E).

**FIGURE 2.**
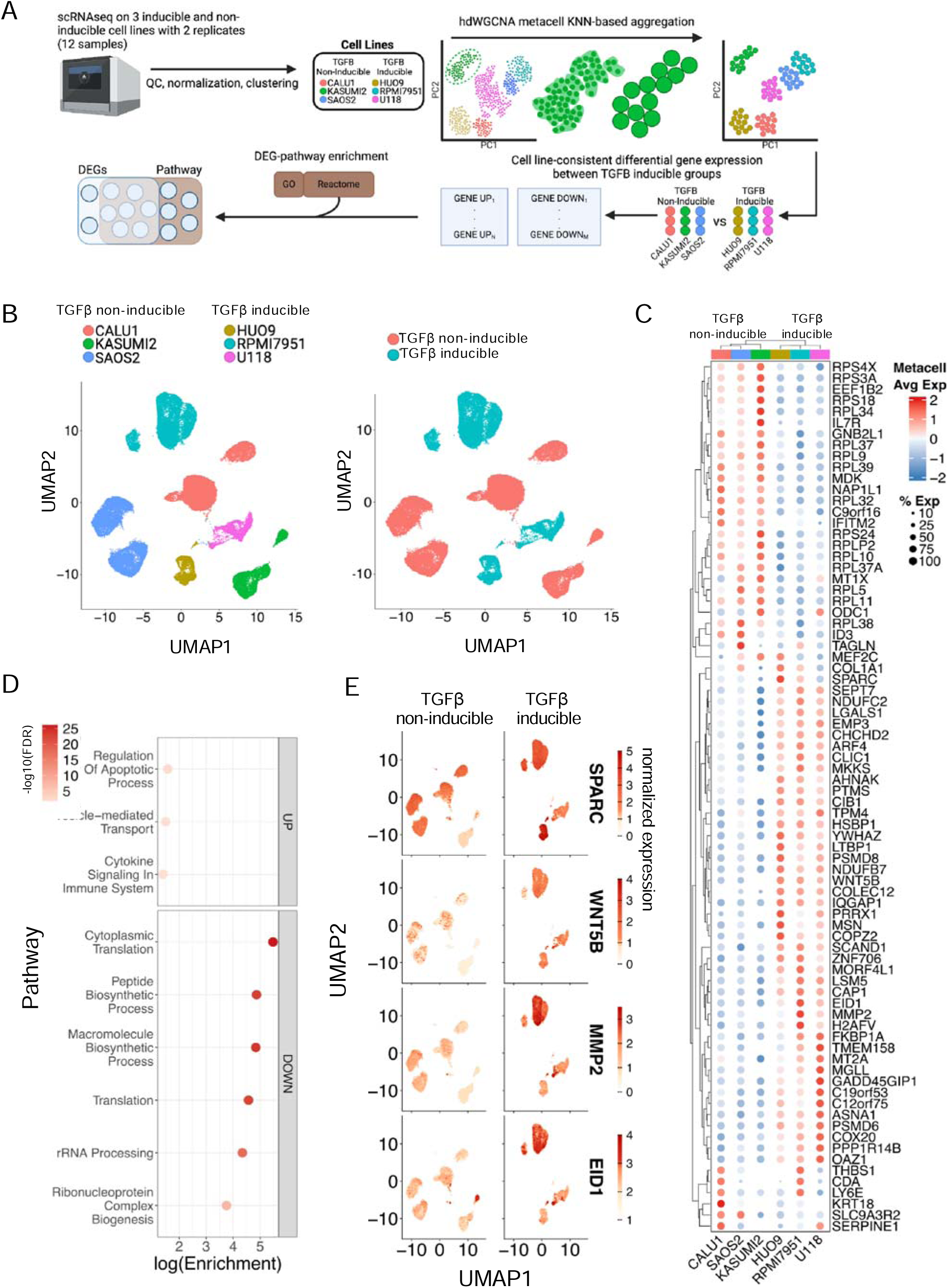
ScRNAseq reveals distinct transcriptional signatures in TGFβ-inducible cancer cell lines. **(A)** Schematic overview of the scRNAseq workflow. TGFβ-inducible (HUO9, RPMI7951, U118) and TGFβ non-inducible (CALU1, KASUMI2, SAOS2) cell lines were sequenced after TGFβ stimulation. Gene expression data were subjected to quality control, normalization, and clustering, followed by hdWGCNA metacell aggregation. DEGs between TGFβ-inducible and non-inducible cell lines were identified and enriched for pathways in Gene Ontology (GO) and Reactome databases using hypergeometric tests implemented in Enrichr. **(B)** UMAP visualization of scRNAseq data from TGFβ-inducible (blue) and non-inducible (red) cell lines. Cell lines did not cluster by TGFβ-inducibility. **(C)** Heatmap of metacell-level expression of large-effect DEGs (|log_2_FC| > 0.5) across TGFβ-inducible (blue) and non-inducible (red) cell lines, represented by average expression (high = red, low = blue) and percent metacell expression by dot size. **(D)** Pathway enrichment analysis of DEGs between TGFβ-inducible and non-inducible cells. Upregulated DEGs were enriched in apoptotic processes and cytokine signaling, while downregulated genes were associated with translation and ribosomal function. **(E)** UMAP plots showing single-cell expression patterns of key upregulated genes in TGFβ-inducible cells, including *SPARC, WNT5B, MMP2*, and *EID1*. These genes demonstrated a marked increase in expression in TGFβ-inducible cells compared to non-inducible cells, consistent with metacell analysis.

### Network analysis of TGF**β**-responsive transcriptional signatures

To further elucidate molecular mechanisms involved in the TGFβ-LRRC15 axis, and to better understand the biological role of the DEGs within LRRC15’s regulation, we employed scRNAseq-based gene network modeling approaches to capture both gene regulatory networks (GRNs) and gene coexpression signatures that distinguished the TGFβ-responsive cell lines (Figure 3A). GRNs consist of nodes and edges representing genes and regulatory interactions, respectively. To construct a GRN for the TGFβ-LRRC15 axis, we used SCING (21), a bagging gradient-boosting machine learning approach to predict regulatory relationships between genes. This method was previously shown to effectively predict gene perturbation effects and uncover key driver genes for disease (21). We incorporated all samples to build an aggregate GRN and identified 41 highly connected SCING GRN modules through Leiden clustering (22). We also constructed a gene co-expression network across samples using hdWCGNA based on gene correlation (19), identifying 25 co-expression modules. The expression levels of individual gene regulatory or coexpression modules from both SCING and hdWCGNA networks were computed for each cell and subsequently for each metacell using hdWGCNA’s module eigengene calculation (19). Modules associated with TGFβ inducibility were identified (Figure 3A).

**FIGURE 3.**
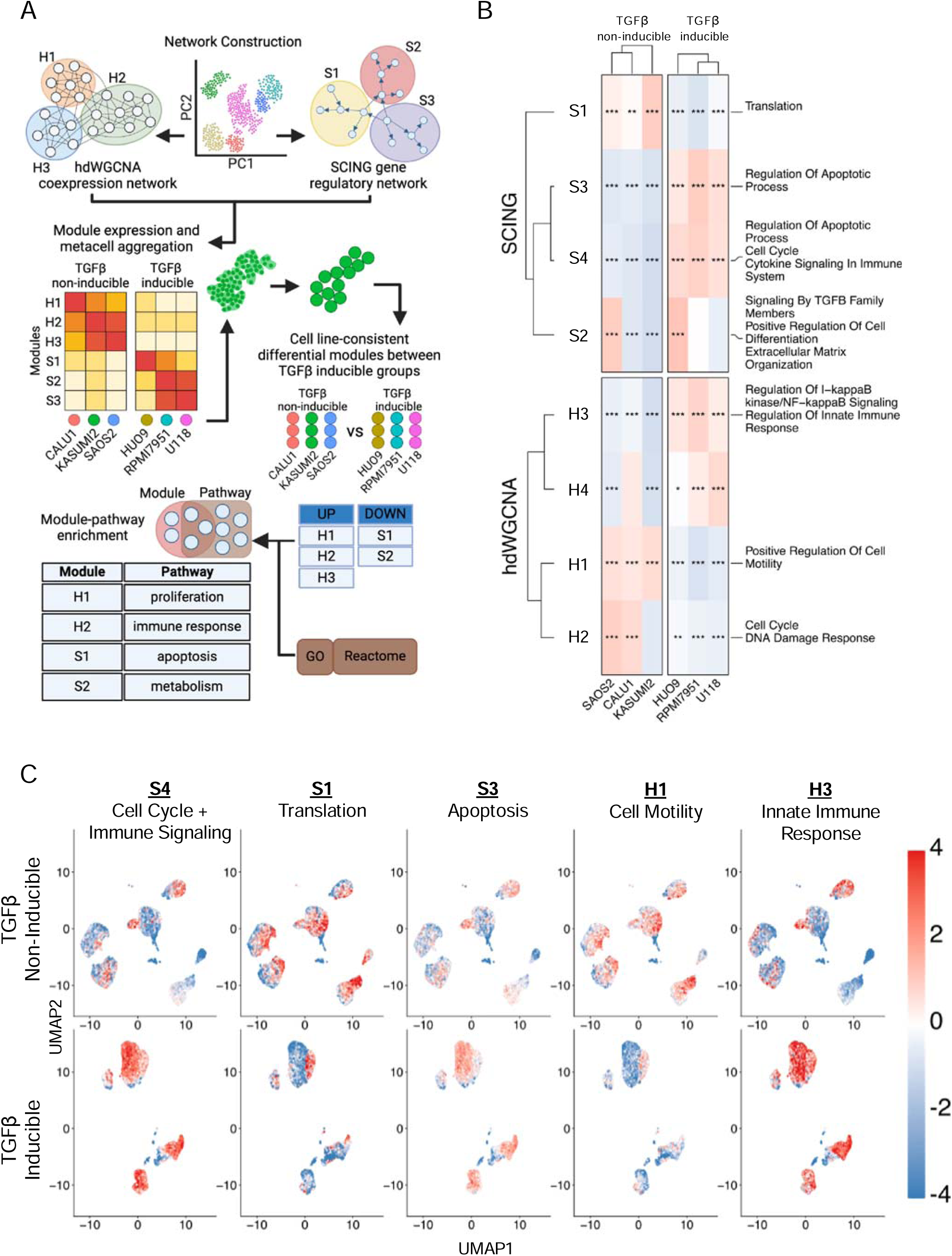
Network analysis of TGFβ-responsive transcriptional signatures. **(A)** SCING and hdWGCNA were used to model gene regulatory networks and coexpression networks, respectively, in TGFβ-responsive and non-responsive cell lines. SCING GRNs were constructed to predict regulatory relationships between genes and identify SCING modules containing genes with regulatory relationship. hdWGCNA co-expression networks were built to capture gene coexpression relations and identify coexpression modules. Expression patterns of SCING and hdWGCNA modules in metacells were compared between TGFβ-inducible and non-inducible cell lines to identify differential modules. **(B)** Heatmap showing expression patterns in SCING and hdWGCNA network modules across cell lines. Modules upregulated and associated with TGFβ responsiveness (in red) included pathways related to cell cycle, cytokine signaling, immune response, and apoptosis. **(C)** UMAP plots displaying single-cell expression patterns of key biological processes of network modules in TGFβ-inducible (top) and non-inducible (bottom) cell lines, confirming the patterns seen in **(B)**.

We identified differential expression in SCING modules S1, S3, and S4, and hdWGCNA modules H1 and H3 (Figure 3B) that consistently differentiated TGFβ-inducible from non-inducible cell lines, and 11 other modules with consensus across 5 cell lines (Supplemental Figure 2A). Gene Ontology and Reactome pathway analysis revealed upregulation of biological pathways related to apoptotic processes and cytokine and immune signaling, with a downregulation of translation (consistent with DEG analysis in Figure 2D), cell motility regulation and DNA damage response in TGFβ-inducible cell lines. UMAP visualization of the cellular expression of the modules and their associated pathways confirmed these patterns (Figure 3C). Surprisingly, cell motility pathways were also downregulated in TGFβ-inducible cell lines (Figure 3C), which contrasted LRRC15’s proposed role as a driver of invasiveness and cell migration in tumor cells.

### Integrated high-throughput drug screening and scRNAseq data reveals converging molecular pathways involved in LRRC15’s regulation

After identifying modules and DEGs associated with TGFβ inducibility from scRNAseq analysis, we combined these results to carry out an *in-silico* drug screen to identify the top candidate drugs and genes within the TGFβ-LRRC15 axis (Figure 4A) and further integrated the results with those from the *in-vitro* drug screen (Figure 1). The L1000 database is a curated library of thousands of compounds and their corresponding affected genes, based on *in-vitro* screening assays across different organisms, tissues, drug dosages, and timepoints (23). After filtering the database for all human drug signatures, we performed a Fisher’s exact test against hdWGCNA– and SCING-derived network modules, all significant (adjusted p-value < 0.05) DEGs, and all large-effect DEGs (|log_2_FC| > 0.5) to identify L1000 compounds that, upon treatment, displayed gene signatures that overlapped with our scRNAseq findings (Figure 4A). This *in-silico* drug screening identified 26 compounds whose gene signatures matched our scRNAseq results, and that were also present in our *in-vitro* drug screen (Figure 1D). We also identified 14 gene targets shared across all analyses (Figure 4B, Table 1). Visualizing these gene targets around *TGF*β*1* and *LRRC15* in the SCING GRN revealed interconnected subnetworks with potential regulators of the TGFβ-LRRC15 axis (Figure 4C).

**FIGURE 4.**
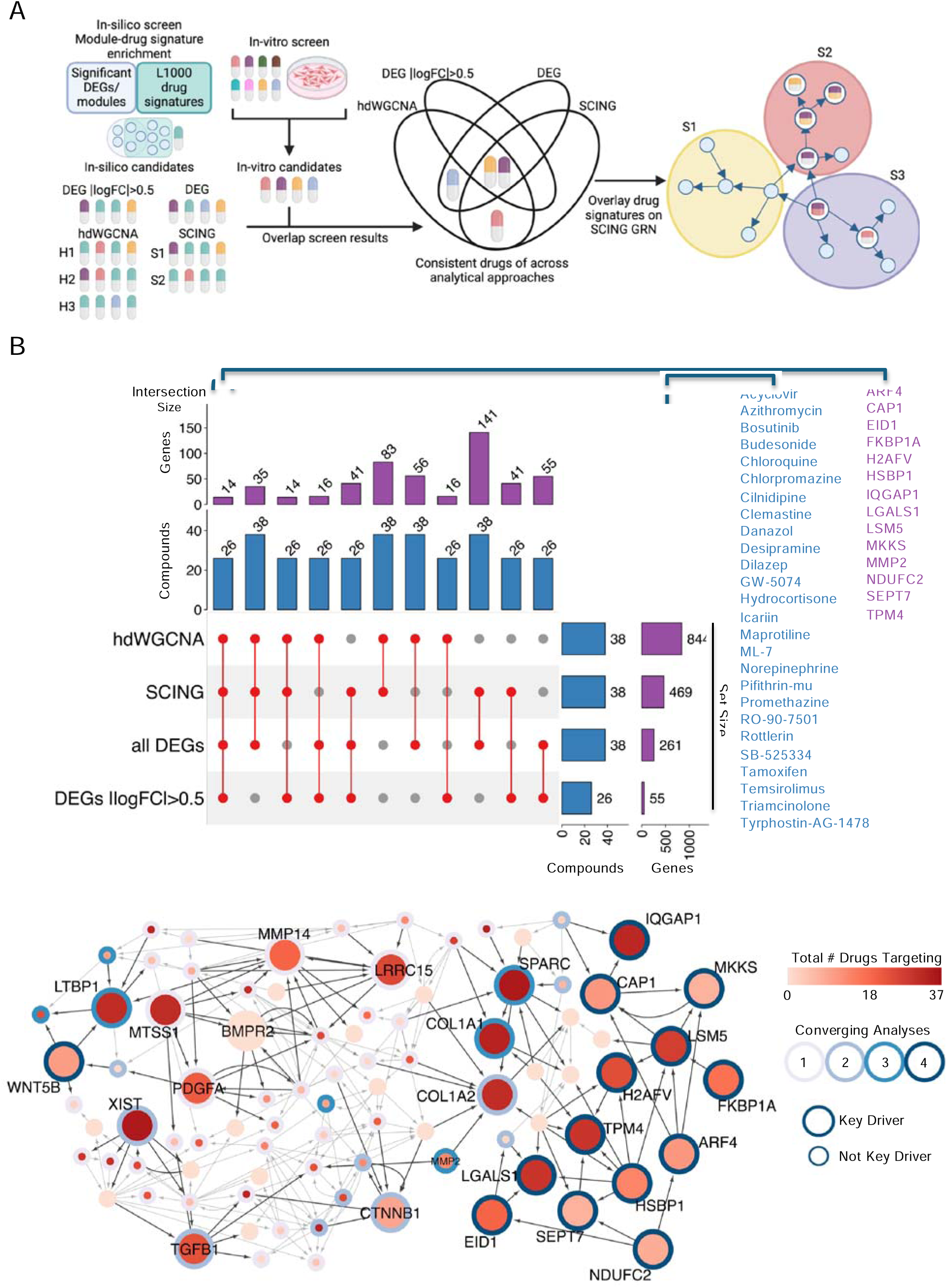
Integration of high-throughput drug screening and scRNAseq reveals converging molecular pathways regulating LRRC15 in TGFβ-inducible cells. **(A)** *In-silico* and *in-vitro* drug screening approaches were combined to identify candidate compounds targeting the TGFβ-LRRC15 axis. The *in-silico* screen utilized the L1000 drug database to map compounds against DEGs and differential hdWGCNA and SCING modules, identifying drug and corresponding genes signatures consistent across analyses. Overlap between the *in-silico* and *in-vitro* drug screens against scRNAseq results highlighted 26 compounds as top candidates for targeting LRRC15 regulation. **(B)** Illustration of the overlap between hdWGCNA, SCING, and DEG datasets, highlighting 26 candidate compounds hits from screening (Figure 1) and 14 gene targets shared across analyses. **(C)** Visualization of gene targets in a gene regulatory network. Key driver analysis revealed potential regulators (larger nodes) of the TGFβ-LRRC15 pathway. Circle fill indicates the total number of compounds targeting each gene, and the number of overlapping analyses identifying the gene as significant is denoted by outline color intensity.

**TABLE 1.** Top gene candidates for modulating the TGFβ-LRRC15 axis.

| Gene | Average Expression Log2 Fold Change | Significant Modules | Compounds |
| --- | --- | --- | --- |
| LGALS1 | 2.63 | S3, S4, H1, ALL_DEG, DEG_0.5 | Danazol, GW-5074, Hydrocortisone, RO-90-7501, Rottlerin, Bosutinib, Budesonide, Cilnidipine, Dilazep, Perphenazine, Pifithrin-mu, Pimozide, SB-525334, Tamoxifen, Temsirolimus, Tyrphostin-AG-1478, Chlorpromazine, Clemastine, Desipramine, Triamcinolone, Acyclovir, Azithromycin, BW-723C86, Chloroquine, Clomipramine, Maprotiline, Mibefradil, Norepinephrine, TTNPB |
| SPARC | 1.96 | S4, ALL_DEG, DEG_0.5 | Bosutinib, Budesonide, Chlorpromazine, Danazol, Desipramine, GW-5074, Hydrocortisone, Maprotiline, Promethazine, Rottlerin, SB-525334, Tamoxifen, Temsirolimus, Triamcinolone, Tyrphostin-AG-1478, Acyclovir, Bexarotene, Chloroquine, Cilnidipine, Clemastine, Desmethylozapine, Dilazep, Mibefradil, Perphenazine, RO-90-7501, TTNPB, Arvanil, Azithromycin, BW-723C86, Clomipramine, Flupentixol, Icarin, Pifithrin-mu, Pimozide, Tenidap |
| FKBP1A | 1.74 | S3, S4, H1, ALL_DEG, DEG_0.5 | Tyrphostin-AG-1478, Chlorpromazine, Danazol, Hydrocortisone, Budesonide, Chloroquine, Mibefradil, Tamoxifen, Temsirolimus, TTNPB, Cilnidipine, GW-5074, Pifithrin-mu, Pimozide, Rottlerin, Tenidap, Triamcinolone |
| TPM4 | 1.51 | S3, S4, H1, ALL_DEG, DEG_0.5 | Chlorpromazine, Promethazine, RO-90-7501, Rottlerin, Budesonide, Dilazep, Hydrocortisone, Tamoxifen, Temsirolimus, Tyrphostin-AG-1478, Acyclovir, Azithromycin, Bosutinib, Cilnidipine, GW-5074, ML-7, Norepinephrine, Perphenazine, Pimozide, Triamcinolone, BW-723C86, Chloroquine, Clomipramine, Danazol, Flupentixol, Maprotiline, Pifithrin-mu, SB-525334, TTNPB |
| MMP2 | 1.48 | H1, ALL_DEG, DEG_0.5 | Cilnidipine, Desipramine, GW-5074, Norepinephrine, Perphenazine, Clemastine, Hydrocortisone, Maprotiline, Pifithrin-mu, Pimozide, RO-90-7501, Triamcinolone, Tyrphostin-AG-1478 |
| HSBP1 | 1.24 | S3, S4, H1, ALL_DEG, DEG_0.5 | Promethazine, GW-5074, Rottlerin, Tamoxifen, Bexarotene, Hydrocortisone, Azithromycin, Budesonide, Chloroquine, Chlorpromazine, Cilnidipine, ML-7, Pifithrin-mu, Triamcinolone |
| *H2AFV* | 1.13 | S3, S4, H1, ALL_DEG,<br>DEG_0.5 | Danazol, Hydrocortisone, Promethazine, Rottlerin, Budesonide, Chlorpromazine, Desipramine, Dilazep, GW-5074, Icarin, Perphenazine, Tamoxifen, Tyrphostin-AG-1478, Acyclovir, Cilnidipine, Clomipramine, RO-90-7501, SB-525334, Triamcinolone, Chloroquine, Mibefradil, Pimozide, Tenidap, TTNPB |
| *ARF4* | 1.04 | S3, S4, H1, ALL_DEG,<br>DEG_0.5 | Danazol, Tyrphostin-AG-1478, Chlorpromazine, Cilnidipine, Rottlerin, Chloroquine, GW-5074, Hydrocortisone, Mibefradil, Pimozide, Tamoxifen |
| *LSM5* | 1.01 | S3, S4, H1, ALL_DEG,<br>DEG_0.5 | Chlorpromazine, Danazol, GW-5074, RO-90-7501, Rottlerin, Bosutinib, Budesonide, Cilnidipine, Desipramine, Hydrocortisone, Perphenazine, Pifithrin-mu, Pimozide, Temsirolimus, Acyclovir, Chloroquine, Clemastine, Dilazep, Norepinephrine, Tamoxifen, TTNPB, Tyrphostin-AG-1478, Arvanil, BW-723C86, Maprotiline, SB-525334, Triamcinolone |
| *EID1* | 0.97 | S3, S4, H1, ALL_DEG,<br>DEG_0.5 | Cilnidipine, Budesonide, Desipramine, GW-5074, Hydrocortisone, Perphenazine, RO-90-7501, TTNPB, Acyclovir, Arvanil, Azithromycin, Bosutinib, Chlorpromazine, Clemastine, Desmethylozapine, Rottlerin, SB-525334, Tamoxifen, Triamcinolone |
| *SEPT7* | 0.9 | S3, S4, H1, ALL_DEG,<br>DEG_0.5 | Danazol, Desipramine, Pimozide, GW-5074, Tyrphostin-AG-1478, Bosutinib, Chloroquine |
| *MKKS* | 0.87 | S3, S4, H1, ALL_DEG,<br>DEG_0.5 | Danazol, Maprotiline, Chlorpromazine, Rottlerin, Tyrphostin-AG-1478, Mibefradil, Tenidap |
| *NDUFC2* | 0.82 | S3, S4, H1, ALL_DEG,<br>DEG_0.5 | Budesonide, Danazol, GW-5074, Bosutinib, SB-525334, Pimozide, Triamcinolone, TTNPB |
| *CAP1* | 0.77 | S3, S4, H1, ALL_DEG,<br>DEG_0.5 | GW-5074, Promethazine, Budesonide, Rottlerin, Bosutinib, Tamoxifen, Maprotiline, Pifithrin-mu, Pimozide, Triamcinolone, Tyrphostin-AG-1478 |
| *IQGAP1* | 0.64 | S3, S4, H1, ALL_DEG,<br>DEG_0.5 | GW-5074, Hydrocortisone, RO-90-7501, Rottlerin, Tamoxifen, Tyrphostin-AG-1478, Budesonide, Chlorpromazine, Norepinephrine, Perphenazine, Pimozide, Temsirolimus, Acyclovir, Azithromycin, Chloroquine, Cilnidipine, Clemastine, Clomipramine, Desmethylozapine, Dilazep, Mibefradil, SB-525334, TTNPB, Arvanil, Bosutinib, BW-723C86, Desipramine, Flupentixol, Icarin, Maprotiline, Tenidap, Triamcinolone |
| *WNT5B* | 0.6 | S2, H1, ALL_DEG, DEG_0.5 | Budesonide, Tamoxifen, Temsirolimus, Bosutinib, Chloroquine, GW-5074, Hydrocortisone, Pifithrin-mu, RO-90-7501, Rottlerin |

### TGF**β**-induced LRRC15 expression can be modulated by siRNA knockdown of candidate genes

To test the functional relevance of genes identified in overlapping analyses (Figure 4) within the TGFβ-LRRC15 pathway, candidate genes were knocked down in TGFβ-responsive cell lines via small interfering RNA (siRNA). Following a similar protocol to our compound screening approach (Figure 1), cells were incubated in low-serum media before transfection with the siRNA construct. After transfection, cells were treated with TGFβ and assessed for LRRC15 expression via confocal microscopy. As a positive control, we validated siRNA transfection and knockdown using three LRRC15-targeting siRNAs (Figure 5A). Transfection with siLRRC15 reduced LRRC15 protein levels by an average of 98.61 ± 0.47% in untreated cells, and 94.21 ± 2.02% in TGFβ-treated cells. As an additional control in subsequent siRNA experiments, we also utilized siRNAs targeting *TGF*β*R2*, given the receptor’s established role in LRRC15 CAFs (13).

**FIGURE 5.**
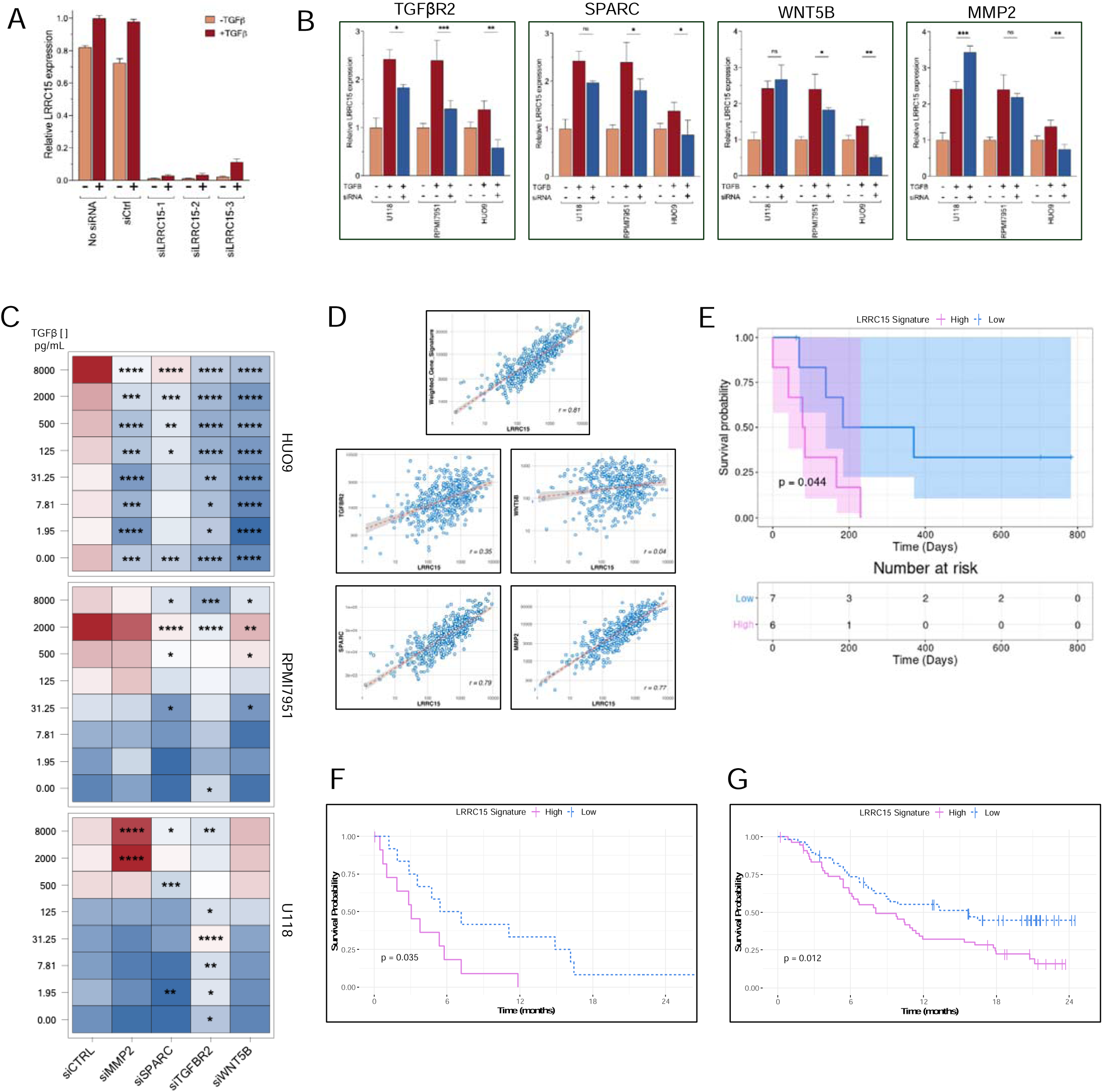
High expression of LRRC15 activating genes predicts immunotherapy resistance and tumor progression. **(A)** Control experiments using LRRC15-targeted siRNA in TGFβ-responsive cancer cell lines significantly reduced LRRC15 expression, overcoming TGFβ-induced upregulation (pC<C0.0005). **(B)** Response to siRNA knockdown of target genes (n=3 per condition) identified within the TGFβ-LRRC15 axis from integrated analysis of scRNAseq and compound screening is not consistent across cell lines, highlighting the importance of genetic background within the TGFB-LRRC15 pathway. **(C)** Heatmap of LRRC15 staining intensity after siRNA gene knockdown across increasing TGFβ concentrations. Significant reductions in LRRC15 expression were observed for *TGF*β*R2, SPARC, WNT5B,* and *MMP2* in osteosarcoma (HUO9), while effects gene knockdown in melanoma (RPMI7951), and glioblastoma (U118) cells were less ubiquitous. **(D)** Correlation plots of LUSC tumor expression from the TCGA database reveal a strong correlation between *LRRC15* and *MMP2* (r = 0.77) and *SPARC* (r = 0.79), but weak correlations between *LRRC15*, *WNT5B*, and *TGFBR2*. There is also a strong correlation between the four genes and *LRRC15* (r = 0.81). **(E)** Kaplan-Meier survival curve demonstrating the predictive power of the 5 genes (including *LRRC15*) in LUSC patients treated with anti-PD1 immunotherapy. Patients stratified by high (red) vs. low (blue) gene expression had significantly different survival outcomes (p =C0.044). **(F)** Kaplan-Meier survival analysis in metastatic clear cell renal cell carcinoma patients receiving anti-PD1 therapy. The *LRRC15*-related genes significantly predicted 2-year survival outcomes in ccRCC patients (pC=C0.035). **(G)** Finally, in metastatic bladder cancer patients receiving atezolizumab (anti-PD1) immunotherapy, high expression of *LRRC15*-related genes was significantly predictive of poorer prognosis in immune-excluded tumors (pC=C0.012), with no significant predictive power in immune-infiltrated or desert tumors (Supplemental Figure 8B).

We then tested siRNAs targeting various genes identified as potential nodes within the TGFβ-LRRC15 axis, including *EID1*, *H2AFV*, *IQGAP1*, *LGALS1*, *MMP2*, *SPARC*, *TGF*β*R2*, *TPM4*, and *WNT5B* (Figure 5B). Among these, *EID1* (EP300 interacting inhibitor of differentiation 1), significantly induced LRRC15 expression upon *EID1* knockdown (p = 0.0059). As a known inhibitor of p300-mediated transcription and differentiation in fibroblast cell populations (24), the inhibition of *EID1* may lead to upregulated p300 activity, possibly explaining the increased LRRC15 expression observed in our studies. While this finding presented an intriguing avenue for further exploration, our primary focus remained on genes whose knockdown led to reduced LRRC15 expression. In addition, we sought to focus solely on genes that were co-expressed with *LRRC15* to improve the feasibility of targeting the TGFβ-LRRC15 pharmacologically. Of the genes tested, *MMP2*, *SPARC* and *WNT5B*, as well as *TGF*β*R2,* significantly impacted TGFβ-induced LRRC15 expression in two or more cell lines, with *TGF*β*R2* and *SPARC* knockdown significantly reducing LRRC15 expression across multiple concentrations of TGFβ treatment (Figure 5B, C). These results support the regulatory role of 4 out of the 9 tested candidate genes in activating the TGFβ-LRRC15 axis and promoting *LRRC15* expression. Notably, the genes that were experimentally validated tended to be the ones with outgoing edges to other genes in the network (Figure 5C); outgoing network edges indicate that the gene likely functions as an upstream regulator.

### Activators of LRRC15 expression correlate to immunotherapy response and tumor progression

The four genes whose knockdown consistently inhibited LRRC15 expression (*MMP2, SPARC, TGF*β*R2, WNT5B*), along with *LRRC15*, became the basis for further exploration into clinical tumor databases. Given the significant impacts of knockdown on TGFβ-induced LRRC15, we reasoned that high expression of these five genes indicated an active TGFβ-LRRC15 axis. We first performed an independent prognostic analysis of the LRRC15 activating genes in cohorts of patients from different tumor entities including breast cancer, glioblastoma, and skin cutaneous melanoma to better understand how our tumor models translated to clinical analyses. Of the three tumor types, our signature acted as a prognostic marker only in breast cancer patients (HR: 2.50, 95% CI 1.66-3.77, p = < 0.001) (Supplemental Figure 6). We instead explored The Cancer Genome Atlas (TCGA) to analyze each gene’s correlation to *LRRC15* across TCGA tumor types to better understand whether *LRRC15* and the potential activators of *LRRC15* were co-expressed in patient tumors (25).

Of the TCGA datasets, *LRRC15* was highly correlated with *MMP2* (r = 0.77, p < 0.0005) and *SPARC* (r = 0.79, p < 0.0005), and moderately correlated with *TGF*β*R2* (r = 0.35, p < 0.0005) in lung squamous cell carcinoma (LUSC) (Figure 5D). Given the strong correlation of 3 out of 4 genes within LUSC tumors, along with LRRC15’s known role in anti-PD1 immunotherapy resistance, we applied our signature to an anti-PD1 immunotherapy trial involving LUSC patients to evaluate whether it could predict immunotherapy resistance. When patients were stratified based on median expression of the 5 genes, we found that *LRRC15* plus the four *LRRC15* activating genes significantly predicted survival outcomes in LUSC (p = 0.044, 13 patients) (Figure 5E). To validate these findings further, we investigated the role of these genes in additional anti-PD1 patient cohorts across other tumor types. We next tested our LRRC15 expression activators in a ccRCC anti-PD1 cohort. Similarly, high expression of *WNT5B, TGF*β*R2, SPARC, MMP2*, and *LRRC15* significantly predicted 2-year survival outcomes (p = 0.035, 24 patients). Notably, when *LRRC15* was removed from analysis, the remaining four genes (*MMP2, SPARC, WNT5B, TGF*β*R2*) failed to predict outcomes (Supplemental Figure 8). However, using *LRRC15* alone did not have the predictive power (Supplemental Figure 8A), emphasizing the importance of the cooperation and co-expression of the 5 genes in contributing to prognostic accuracy.

We evaluated a third cohort from a metastatic bladder cancer trial where patients received atezolizumab (anti-PD1) immunotherapy and were profiled for immune phenotypes based on CD8+ T-cell infiltration within the tumor and surrounding peritumoral region (26). Before undergoing treatment, patients were categorized as “infiltrated” (CD8+ T-cells present within the tumor), “excluded” (CD8+ T-cells confined to the tumor stroma), or “desert” (absence of CD8+ T-cells). Given previous studies implicating LRRC15+ CAFs in T-cell exclusion and exhaustion, we tested the predictive power of the LRRC15-activating genes using transcriptomic data from immune-excluded tumors prior to anti-PD1 therapy. Our analysis demonstrated that high expression of the 5 genes was significantly predictive of patient prognosis (p = 0.012, 113 patients) in immune-excluded tumors (Figure 5F). In contrast, gene expression showed no significant predictive power in tumors classified as immune-infiltrated or desert, or with *LRRC15* alone (Supplemental Figure 7B, 8C). These findings indicate that expression of *MMP2, SPARC, WNT5B* and *TGF*β*R2,* alongside *LRRC15,* can predict patient prognosis and overall response to immunotherapy in immune-excluded tumors, and can accurately reflect the pathobiological consequences of TGFβ-LRRC15 axis signaling. While further analysis is needed, it is likely that immune-excluded tumors harbor large populations of LRRC15+ CAFs and tumor cells, which may contribute to the poor therapeutic response observed by creating a physical barrier as well as imposing immunomodulatory effects. Together, these findings underscore the critical role of LRRC15, along with several key genes within the TGFβ-LRRC15 axis, in shaping the immune-excluded tumor microenvironment. This gene set not only offers a potential biomarker for predicting immunotherapy resistance but also highlights novel therapeutic avenues for targeting LRRC15+ cell populations in resistant cancers.

## DISCUSSION

Deciphering the intricate molecular pathways, genomic determinants, and tumor microenvironmental factors that regulate biomarker dynamics is pivotal for refining targeted diagnostic and therapeutic approaches in oncology. Detailed knowledge of these factors enables the stratification of patient populations most likely to respond to specific treatments, thereby enhancing clinical efficacy (27–28). The absence of pathobiological understanding is a major contributor to the substantial attrition rates in clinical oncology trials, where nearly half of failures are attributed to lack of established relevance of the target to the disease phenotype (27). With the surge in oncology target and biomarker discovery, the translation of these findings into clinically beneficial entities remains elusive, highlighting the critical need for comprehensive molecular profiling to fully realize the promise of personalized medicine in cancer care.

The initial identification of LRRC15’s regulation by TGFβ in MSCs under prolonged supraphysiological cytokine levels (16) prompted a reevaluation of its translational relevance. To address this, our study aimed to investigate LRRC15 regulation under shorter durations and physiologically relevant concentrations of TGFβ. As tumors progress, CAFs and cancer cells increasingly exhibit similar behaviors and interactions, enhancing tumor complexity and resilience. Both cell types are responsive to extracellular molecules such as growth factors and cytokines and are subject to frequent genetic alterations, with cancer cells commonly inducing expression changes in CAFs via paracrine signaling, which can lead to selective CAF expansion (29–30). Furthermore, CAFs maintain tumor-promoting properties independent of direct interactions with cancer cells (31). This convergence of characteristics guided our focus to LRRC15+ MSC-derived cancer cells for deeper insights.

Our investigation revealed two distinct cellular responses to TGFβ following cytokine withdrawal: one subset of cell lines demonstrated LRRC15 upregulation, suggesting an adaptive mechanism, while another remained unaffected. Integrative analysis using scRNAseq, chemical compound screening, and siRNA knockdown pinpointed *TGF*β*R2*, *WNT5B*, *SPARC*, and *MMP2* as key mediators inducing TGFβ-LRRC15 activity. In patient tissue databases, the impact of these genes on TGFβ-LRRC15 activity was observed across a wide range of solid tumors (Supplemental Figure 5), including those demonstrating expression exclusively in CAFs but not in cancer cells. Moreover, a significant correlation emerged with a previously identified *LRRC15*-containing 11-gene panel (Supplemental Figure 9), which was obtained by screening of CAFs in TGFβ-driven tumor microenvironments in PDAC patients (14).

*LRRC15*, along with the four genes regulating its expression, differentiated survival rates and disease progression in tumor settings that were characterized by the exclusion of CD8+ T cells and resistance to ICTs. However, we did not find significant prognostic value in tumors categorized as immune-infiltrated or immune-desert. It could be speculated that these variations may be attributed to immunological mechanisms driven by non-LRRC15 related mechanisms. Among the four genes within the TGFβ-LRRC15 axis, *TGF*β*R2* has previously been established as a regulator of LRRC15+ CAF formation in GEMM models (13). Tumor fibrosis is a common consequence of aberrant TGFβ signaling, contributing to immune cell exclusion in the stroma-rich ECM and poor drug delivery (32). Several of these ECM components were significantly co-expressed with LRRC15, such as matrix metalloproteinases (*MMP2, MMP14*) and collagen genes (*COL1A1, COL1A2*). Further, *MMP2* plays a critical role in cancer cells by promoting angiogenesis and cell growth, and by facilitating collagen degradation in the TME, thereby aiding tumor invasion. This highlights its dual function in promoting cancer progression and supporting CAF populations. In parallel, *SPARC*, a secreted protein of the extracellular matrix, has been shown to synergize with TGFβ signaling to boost collagen production and induce canonical TGFβ signaling pathways, such as SMAD2 activation (33). While further research is necessary to fully delineate *SPARC*’s function in tumor tissues, existing studies indicate that its knockdown in pterygium fibroblasts diminishes TGFβ signaling and *MMP2* expression (34), potentially leading to a similar effect on LRRC15+ CAF and cancer cell interactions.

In the current study, we identified a broad spectrum of potential regulatory genes; however, siRNA knockdown of certain genes, such as *H2AFV*, did not significantly affect TGFβ-induced LRRC15 expression in our cancer cell models. The lack of significant effects on TGFβ-induced LRRC15 expression could stem from several factors. First, these genes tend to have more incoming edges (i.e., receivers of regulation by other genes) than outgoing edges (i.e., acting as upstream regulators) in the networks. Second, although *H2AFV* is upregulated in various cancers, it primarily operates through epigenetic mechanisms such as DNA methylation reprogramming (35). The epigenetic influence of H2A and its variants, which often have overlapping functions, adds a layer of complexity to gene regulation (36). Lastly, even with effective siRNA transfection, residual protein expression might maintain *H2AFV*’s role within the TGFβ-LRRC15 axis (37). Further investigations, possibly employing CRISPR/Cas9 gene editing for more complete gene disruption, are necessary to fully understand these genes’ involvement in LRRC15 expression (38).

Our pioneering small molecule screens identified a multitude of compounds capable of disrupting the TGFβ-LRRC15 pathway (Figure 1), potentially providing alternative strategies to overcome the pro-tumorigenic effects of TGFβ without direct inhibition of the pathway itself. This is especially important given the clinical challenges associated with TGFβ receptor inhibitors, where inhibition often leads to adverse off-target effects. For example, bosutinib, a Bcr-Abl tyrosine-kinase inhibitor, is actively studied in chronic myeloid leukemia (CML) as a combination therapy with anti-PD1 immunotherapy (39). In solid tumors, bosutinib may be able to play a similar complementary role to immune checkpoint blockade by inhibition of TGFβ-related immunosuppressive pathways. In addition, the significant downregulation of DNA damage response processes in TGFβ-inducible cell lines may expose a potential therapeutic vulnerability in these cell populations. The positive hits identified from our drug screening, particularly those supported by both *in-vitro* and *in-silico* screens, could be further explored for such applications in the future.

Upon exploring LRRC15 as a biomarker for therapeutic resistance, our analysis of TCGA sarcoma datasets suggested that *LRRC15* alone is insufficient as a prognostic biomarker for aggressive disease and overall survival and is not predictive of therapeutic outcomes in immunotherapy trials (Supplemental Figure 7C). These observations contrast with prior clinical findings in an osteosarcoma cohort where LRRC15 was identified as a prognostic marker for therapy resistance and reduced survival (10). This discrepancy highlights that *LRRC15* expression alone may not solely drive tumor aggressiveness. Integrating the results of this study, we hypothesize that LRRC15 functions primarily as a bystander protein, indicative of an extensive network of TGFβ-driven pathobiological processes prevalent in the microenvironment of aggressive, immunoresistant tumors.

We have previously demonstrated that LRRC15-targeted PET and SPECT imaging is applicable for non-invasive detection, quantification, and monitoring of lesion-specific LRRC15 (15). LRRC15, with its low expression in non-cancerous and healthy tissues and high expression in aggressive tumors, serves as an ideal therapeutic target for RIT. This strategy facilitates the selective delivery of cytotoxic radionuclides to specifically target and ablate both cancer cells and CAFs that contribute to treatment resistance and metastatic growth. In addition to the ablation of LRRC15+ cell populations, tumors treated with ^177^Lu-LRRC15-RIT exhibited downregulation of *SPARC, MMP2, TGF*β*R2*, and *WNT5B* (Supplemental Figure 4). These findings partially confirm that *LRRC15* is co-expressed with other crucial factors involved in driving tumor progression and resistance.

Finally, with the integration of next-generation sequencing and comprehensive genomic profiling in clinical decision making, it has become possible to stratify patients more accurately based on tumor molecular profiles (40). Large-scale studies like NCI-MATCH have successfully matched patients to treatment options, with 38% of patients having tumor molecular profiles that were actionable by currently available clinical trials (41–42). Other molecular sequencing studies such as ComboMATCH have expanded beyond simple biomarker analysis to incorporate circulating tumor DNA, histology, and RNA sequencing of patient tumors (43), highlighting the feasibility of applying assessment of LRRC15-related profiles to guide decision-making and individualized treatment in a clinical setting.

In summary, by computationally integrating high-throughput small molecule screening data with scRNA sequencing, we identified *TGF*β*R2, MMP2, SPARC*, and *WNT5B* as key mediators driving LRRC15 upregulation in response to TGFβ signaling. These molecules, secreted by both cancer cells and fibroblasts, are critical in promoting EMT, during which LRRC15 expression is significantly elevated. Their coordinated activity shapes a fibrotic and immunosuppressive tumor microenvironment, predicting resistance to immune checkpoint inhibitors and correlating with aggressive cancer phenotypes. These findings elucidate the molecular architecture of the TGFβ-LRRC15 axis, revealing a cascade of effector molecules that present potential targets for the development of inhibitors and modulators to disrupt these pathways. This improved understanding may also provide insight into the mechanisms underlying immunoresistance, offering a basis for exploring novel diagnostic and therapeutic strategies to enhance the efficacy of immunotherapy.

## MATERIALS AND METHODS

A detailed description of materials, methods, and equipment from this study is provided in the supplementary materials.

### In vitro evaluation and quantification of LRRC15

Detailed descriptions of cell lines used, as well as LRRC15-based flow cytometry, confocal microscopy, and confocal microscopy-based screening methods are available in the supplementary materials.

### Silencer RNA

Silencer RNAs targeting selected genes were obtained from ThermoFisher Scientific (catalog #4392420). 72 h after siRNA transfection, cells were treated with 0-8 ng/mL TGFβ1 for 24 h, followed by fixation and LRRC15 staining as described above. Successful transfection was confirmed by loss of LRRC15 expression in all three siLRRC15 conditions. Plates were imaged using a ImageXpress Micro Confocal High-Content microscope, and images were analyzed with a custom module in MetaXpress (see supplementary methods). Total LRRC15 expression for each siRNA condition was quantified by measuring the average integrated intensity of LRRC15 staining in each well, normalized to the staining intensity in non-transfected, TGFβ1-treated and untreated controls.

### scRNAseq Analysis

scRNAseq datasets from all cell lines (6 cell lines, 2 replicate per cell line) were analyzed together using Seurat’s library size normalization, log transformation, scaling, principal component analysis (PCA), K-nearest neighbor detection, and UMAP visualizations (44–45). Standard quality control (QC) procedures were applied to filter out poor quality cells, keeping only cells with mitochondrial gene percentage below 5% and gene count between 200 and 5000. We kept genes that were expressed in at least 100 cells. QC plots are reported in Supplemental Figure 3. In total, 57,771 cells passing QC across all cell lines were kept for downstream analysis. With hdWGCNA’s metacell approach (46), cells with similar expression were aggregated and averaged within each cell line to reduce gene expression sparsity for differential gene expression analysis and network modeling. To identify DEGs associated with TGFβ inducibility while considering the sample size (2 replicates per cell line), a one sample-vs-opposite group DEG analysis was performed consisting of 6 separate Wilcoxon rank sum tests comparing each cell line (e.g., one cell line in the TGFβ inducible group) to all cell lines in the opposite comparison group (e.g., all cell lines in the non-inducible group). Only DEGs that were consistent across all comparisons were considered for downstream analysis to avoid cell line-specific variation that could heavily skew the gene counts distribution and produce false positives. All significant DEGs passing false discovery rate (FDR) <5% and large-effect DEGs with an absolute value log fold change of at least 0.5 were used as two DEG sets for downstream analysis.

### Pathway Enrichment Analysis

Pathway enrichment of DEGs and SCING and hdWGCNA modules was conducted using the EnrichR R package for Gene Ontology Biological Process and Reactome databases separately (47). Benjamini Hochberg (BH) correction was performed to control the false discovery rate.

### Network Analysis

SCING regulatory network inference (21) was employed on scRNAseq data, implementing a bootstrapping and gradient boosting strategy to generate and aggregate gene regulatory networks to produce a consensus GRN across subsamples of the data (21). With the final GRN, Leiden clustering was used to identify highly connected subnetworks, or modules, and module expression profiles were generated for each cell line using the module eigengene approach from hdWGCNA (19). hdWGCNA was also run to establish a correlation-based co-expression network method for single cell data (19). Modules from this network were obtained through the method’s topological overlap matrix and corresponding expression using the module eigengene approach (19). We performed module-trait association analysis to correlate TGFβ inducibility with the modules. To identify modules associated with TGFβ inducibility, the one sample-vs-opposite group approach mentioned in the DEG analysis was performed on the module expression data for SCING and hdWGCNA modules separately.

### In Silico Drug Repositioning Analysis Using scRNAseq DEGs and Network Modules

To identify drug candidates matching the TGFβ inducible gene sets from scRNAseq (two DEG sets, one with all significant DEGs and the other with large-effect DEGs (|log_2_FC| >0.5); two differential network module sets from SCING and hdWGCNA), we applied a one-tailed Fisher’s exact test between each of the four TGFβ inducible gene sets and curated drug target gene signatures from the L1000 forward drug signature database (23). We subset the L1000 drug signatures to focus on human datasets. BH FDR correction was applied to the overlap analysis to derive significant overlaps at FDR<5% and drugs with fewer than 5 gene overlaps with the scRNAseq signatures were eliminated from the significant results. Significant drugs from our *in-silico* drug repositioning analysis were overlaps with those from our *in-vitro* high throughput screening to identify consistent drugs. The scRNAseq-based TGFβ inducible gene sets targeted by these replicated drugs were used for candidate regulatory gene prioritization, where gene signatures were overlapped across the four TGFβ inducible gene sets targeted by replicated drugs to identify consistent target genes with potential relationships to the TGFβ-LRRC15 axis.

### Key Driver Analysis to Identify Potential Regulators of TGFβ-LRRC15 Axis

Potential regulators of the TGFβ-LRRC15 axis were identified using Key Driver Analysis (KDA) on the SCING GRN (21). KDA uses the GRN topology to identify hub genes with high connectivity to gene sets of interest. It identifies hub gene neighborhoods enriched for the gene sets through a chi-like statistic, and measures significance of the enrichment based on the null distribution of enrichment scores generated from permuted networks. The significant hub genes are proposed as the key drivers of the gene sets of interest. We performed KDA on SCING GRN modules to detect key drivers within each module.

### Immunotherapy Trial Data Analysis

For analysis of the anti-PD1 trial data in advanced bladder cancer (NCT02951767), gene expression data was obtained from the IMVigor210CoreBiologies package. Tumors were filtered by top 75% LRRC15+ before calculation of a gene expression score of LRRC15 and the four LRRC15-related genes. Scores were computed with the eigenWeightedMean function in the MultiGSEA package in R. Kaplan-Meier survival curves were plotted using the Survminer package with R version 4.4.1 environment. LUSC (GSE93157) and RCC anti-PD1 (14) gene expression data were obtained from repositories (LUSC) or as a normalized expression data matrix (RCC).

## Supporting information

AllModuleGenes

AllHTSResults

## SUPPLEMENTAL MATERIALS AND METHODS

### Cell Lines

CALU1 (non-small-cell lung cancer), KASUMI-2 (leukemia), NCI-H196 (small cell lung cancer), RPMI7951 (melanoma), SAOS2 (osteosarcoma), U118-MG (glioblastoma), U2OS (osteosarcoma) and U87-MG (glioblastoma), were purchased from ATCC. HUO9 (osteosarcoma) was purchased from the Japanese Collection of Research BioSources (Tokyo, Japan). All cell lines were cultured according to the manufacturer’s instructions (base media with 10% FBS) and frequently tested for Mycoplasma by PCR.

### Flow Cytometry

Cells were serum-starved in appropriate base media + 0.3% FBS for 16 hours, followed by treatment with recombinant human TGFβ1 (0.001ng/mL-10ng/mL, Peprotech, #100-21C) for 24 hours. The LRRC15-binding antibody DUNP19 (https://doi.org/10.1101/2024.01.30.577289) was conjugated to Alexa Fluor 647 with an amine-reactive antibody labeling kit (ThermoFisher, #A20186) following manufacturer protocols for labeling. Cells were stained for LRRC15 expression with 100ng/mL DUNP19-AF647 for 45 minutes at room temperature and analyzed using the Attune NxT Flow Cytometer (Invitrogen) in collaboration with UCLA’s Jonsson Comprehensive Cancer Center Flow Cytometry Shared Resource. Data was analyzed using FlowJo (Version 10, BD Biosciences).

### Confocal Microscopy

Cells (0.002 x 10^6^ cells/well) were seeded in appropriate base media + 0.3% FBS for 16 hours in 384-well u-clear flat bottom black plates (Greiner, #781092) before treatment with recombinant human TGFβ1 (0.001ng/mL-10ng/mL) for 24 h. Cells were fixed with 3.7% PFA in PBS for 15 minutes at room temperature, washed, and stained with 100ng/mL DUNP19-AF647 and Hoechst 33342 at a 1:2000 dilution (Invitrogen, #H1399) for 2 hours at room temperature in the dark. Cells were washed with PBS and imaged using a ImageXpress Micro Confocal High-Content Imaging microscope (Molecular Devices). For each well, images were taken at 10x objective with 4 sites imaged per well. For TGFβ inhibition, 0-100uM Galunisertib (MedChemExpress, #HY-13226) was added 16 h prior to addition of 0-10 ng/mL TGFβ1. After 24 h, cells were fixed and stained for LRRC15 as described above.

### Small Molecule Screening

The LOPAC^1280^ compound library (Sigma Aldrich) was used to identify inhibitors that could block TGFβ’s induction of LRRC15 protein expression. Compounds were dissolved at 1 mM in DMSO, and 250 nL of each was transferred into 384-well u-clear flat-bottom black plates, resulting in a final concentration of 5 μM. The liquid transfer was performed using a Biomek automated liquid handler (Beckman Coulter).

TGFβ-responsive cells (0.04 x 10^6^ cells/mL; HUO9, RPMI7951, or U118-MG) were resuspended in media containing 0.3% FBS, transferred to the prepared plates (50 µL/well, 2000 cells), and incubated for 16 h. After incubation, plates were treated with a final concentration of 4 ng/mL TGFβ1 diluted in serum-starved media; this TGFβ1 concentration was chosen because it resulted in maximal LRRC15 induction in flow cytometry and confocal microscopy assays across cell lines. Columns 1-2 of the 384-well plates were treated with TGFβ1 without any compounds, serving as positive controls for LRRC15 induction. In contrast, columns 23-24 were left untreated to establish baseline LRRC15 expression.

After 24 h, cells were fixed with 3.7% PFA in PBS for 15 minutes at room temperature, washed with PBS, and stained with 100 ng/mL DUNP19-AF647 and Hoechst 33342 at a 1:2000 dilution (Invitrogen, #H1399) for 2 h at room temperature in the dark. After staining, cells were washed with PBS before imaging with a ImageXpress Micro Confocal High-Content Imaging microscope (Molecular Devices). For each well, images were taken at 10x objective with 4 sites imaged per well.

### Image Analysis of Confocal Microscopy

Images from the plates were processed using a custom image analysis module in MetaXpress. Nuclei were first identified based on Hoechst 33342 staining, and nuclei count, integrated Hoechst 33342 intensity, and nucleus size were recorded. LRRC15+ cells were defined by staining intensity at or exceeding a threshold of 2000 above background. The number of LRRC15+ cells was normalized to total nuclei count, and the average %LRRC15+ cells per well was calculated by averaging values across all imaged sites within each well.

A standardized Z-score was calculated by subtracting the mean %LRRC15+ value of all wells within the plate and dividing by the standard deviation in %LRRC15+ scoring. Z-scores were similarly calculated for nuclei counts to identify toxic compounds, excluding control wells from all calculations. Hit compounds were defined as any non-toxic compounds (nuclei count Z-score > –1) that reduced LRRC15 staining to a Z-score of < –3.

### scRNAseq Sample Preparation

Single cells were FACS sorted for the top 1% LRRC15+ cells by AlexaFluor647-DUNP19 staining with a target cell population of 10,000 viable cells. Single cells were captured within gel bead emulsions (GEM) according to 10X Genomics’ Chromium Single Cell platform protocols. Within individual GEM droplets, cells are lysed and barcoded before reverse transcription and cDNA pooling. A 3’GEX library was constructed and quality control performed with Agilent ScreenTape Analysis. Libraries were sequenced with Novaseq S4 (2000-2500 M reads per lane), with a read depth of 20,000.

### Network Visualization

Network visualization was performed on the SCING network using Cytoscape (48). The network visualized in Figure 4C include the top drug target candidates from the in-silico drug screen and their direct paths to LRRC15 and TGFB. All neighbors of the drug targets are found in Supplemental Figure 2B. Because these networks focus on the drug targets, some key drivers’ connections are not visualized and thus show little connectivity.

### Silencer RNA

A scramble control, GFP-tagged siRNA, and LRRC15-targeting siRNAs were used as transfection controls. For the siRNA reverse transfection, 1 pmol siRNA was diluted in 5uL Opti-MEM reduced serum media (Gibco, #31985062) and added to a 96-well black uClear plate (Greiner Bio-One, #655090). For each condition, 0.3 uL Lipofectamine RNAiMAX transfection reagent (ThermoFisher Scientific, #13778075) was diluted in Opti-MEM reduced serum media, added to siRNA/Opti-MEM mixture 1:1 v/v, and incubated for 5 m at room temperature. Cells (HUO9, U118-MG, RPMI7951) were harvested and resuspended at 0.025 x 10^6^ cells/mL in antibiotic-free complete media. 100 uL of the cell resuspension was added to each well for a reverse transfection. Transfections were performed in triplicate for each siRNA and control condition.

### Hazard Ratio Analysis

Gene expression data from the different tumor entities were obtained from publicly available repositories for the following entities (breast cancer GSE25066), glioblastoma multiform (PRJNA482620) and skin cutaneous melanoma (phs00452) or in-house generated (pancreatic cancer). The expression data for all genes constituting the LRRC15 signature was extracted. A regression model with time-to-event outcome was performed and the regression coefficients were used to compute the LRRC15 signature as previously described (31699795). Kaplan-Meier survival analysis was performed for the LRRC15 signature for all entities after an optimal-cutoff was determined using the Survminer package with R version 4.4.1 environment.

#### Acknowledgements

This study was supported in part by the UCLA Eli and Edythe Broad Center of Regenerative Medicine and Stem Cell Research Rose Hill Foundation Innovator Award. The study was further supported by NCI R01CA 2010 35, R01CA240711, R01CA229893, DoD W81XWH-18-1-0223, UCLA SPORE in Prostate Cancer (P50 CA092131), JCCC Cancer support grant from NIH P30 CA016042 (PI: Teitell), Knut and Alice Wallenberg Foundation, Bertha Kamprad Foundation, David H. Koch Prostate Cancer Foundation Young Investigator Award, Swedish Research Council, Swedish Cancer Society, SIPEA Foundation, Swedish Childhood Cancer Foundation, John and Augusta Perssons Foundation, Royal Physiographic Society of Lund, Franke and Margareta Bergqvist Foundation, Crafoord Foundation, Lund University Medical Faculty research time allocation award, and IngaBritt and Arne Lundberg Research Foundation. Flow cytometry was performed in the UCLA Jonsson Comprehensive Cancer Center (JCCC) and Center for AIDS Research Flow Cytometry Core Facility that is supported by National Institutes of Health awards P30 CA016042 and 5P30 AI028697, and by the JCCC, the UCLA AIDS Institute, the David Geffen School of Medicine at UCLA, the UCLA Chancellor’s Office, and the UCLA Vice Chancellor’s Office of Research. We also thank the UCLA Technology Center for Genomics and Bioinformatics (TCGB) for preparation of the single cell RNA sequencing, and the UCLA Molecular Screen Shared Resource (MSSR) for their assistance with the compound screening design and implementation. J.T.S. is grateful for support from the German Cancer Consortium (DKTK) and the German Federal Ministry of Education and Research (BMBF; 01KD2206A/SATURN3). We acknowledge support by the Westdeutsche Biobank Essen (WBE, University Hospital Essen, University of Duisburg-Essen, Essen, Germany and the High Throughput Sequencing Unit (Genomics & Proteomics Core Facility, DKFZ) for providing excellent sequencing services. Lastly, we express our sincere gratitude to Sven-Erik Strand for his kind assistance with administrative tasks at Lund University.

## Data Availability

Data available on request from the authors.

## Author Contributions

CMS: writing (original draft), writing (review and editing), data collection, formal analysis

MC: writing (original draft), writing (review and editing), data collection, formal analysis

MA: writing (original draft), writing (review and editing)

JEP: data collection, formal analysis

JT: formal analysis

SSL: formal analysis

LM: data collection, formal analysis

WZ: data collection

CY: data collection

AR: data collection

MTA: formal analysis

KH: writing (review and editing)

SS: formal analysis, writing (review and editing)

BS: formal analysis, writing (review and editing)

KL: writing (review and editing)

JTS: writing (review and editing)

RD: writing (review and editing), formal analysis, conceptualization

XY: writing (original draft), writing (review and editing), conceptualization

DU: writing (original draft), writing (review and editing), conceptualization

## SU

**SUPPLEMENTAL FIGURE 1.**
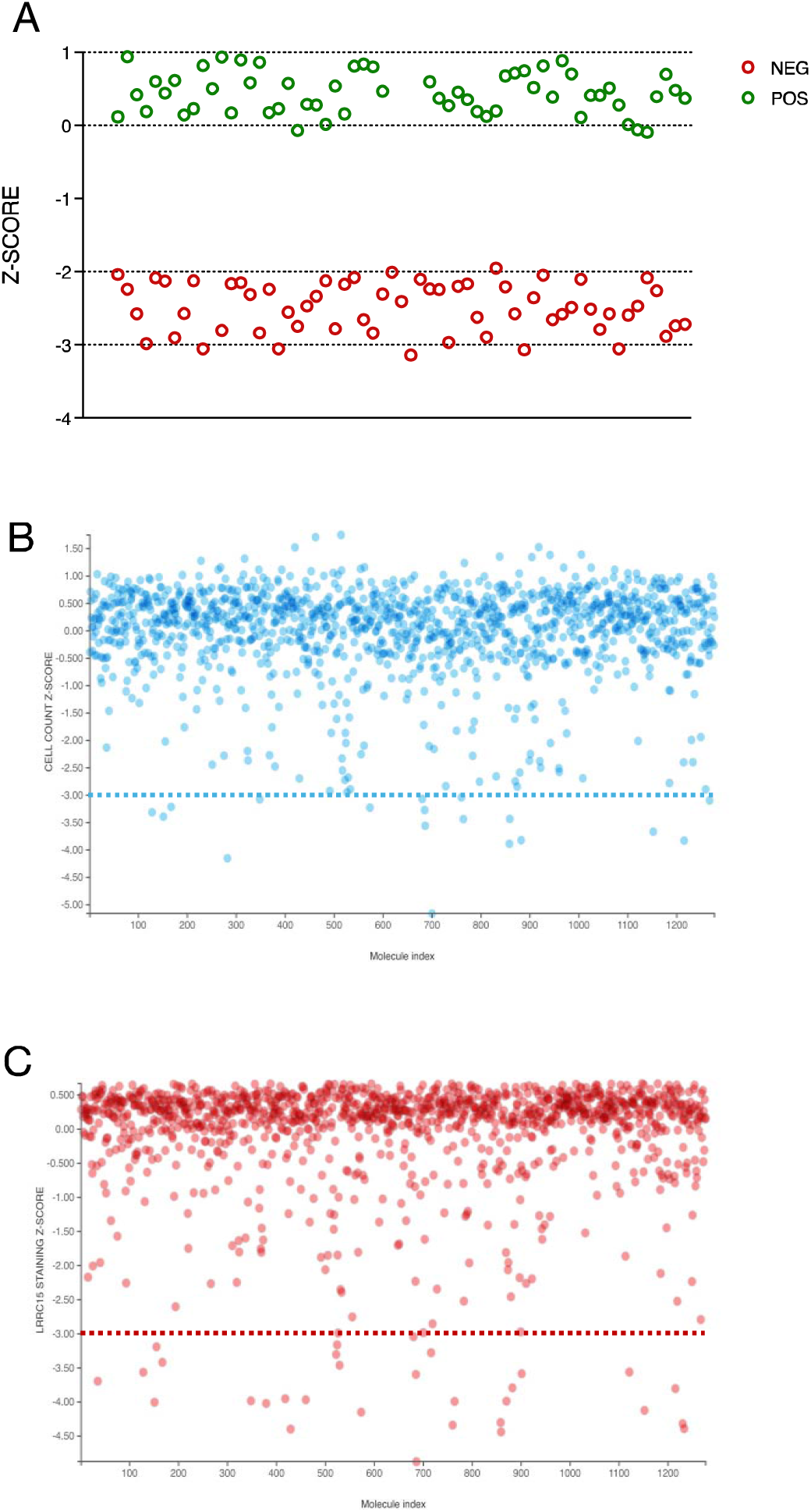
Compound screening controls. **(A)** Summary of TGFβ-treated (POS, green) or non-treated (NEG, red) LRRC15 staining controls, represented by Z-score. **(B)** Cell count per well across 1,280 screened compounds, represented by Z-score. Dotted blue line represents compound toxicity cutoff where compounds were omitted due to cytotoxic effects (Z-score < –3). **(C)** LRRC15 staining per well across 1,280 screened compounds, represented by Z-score. Dotted red line represents a LRRC15 staining Z-score of < –3, which was determined to be a hit within compound screens.

**SUPPLEMENTAL FIGURE 2.**
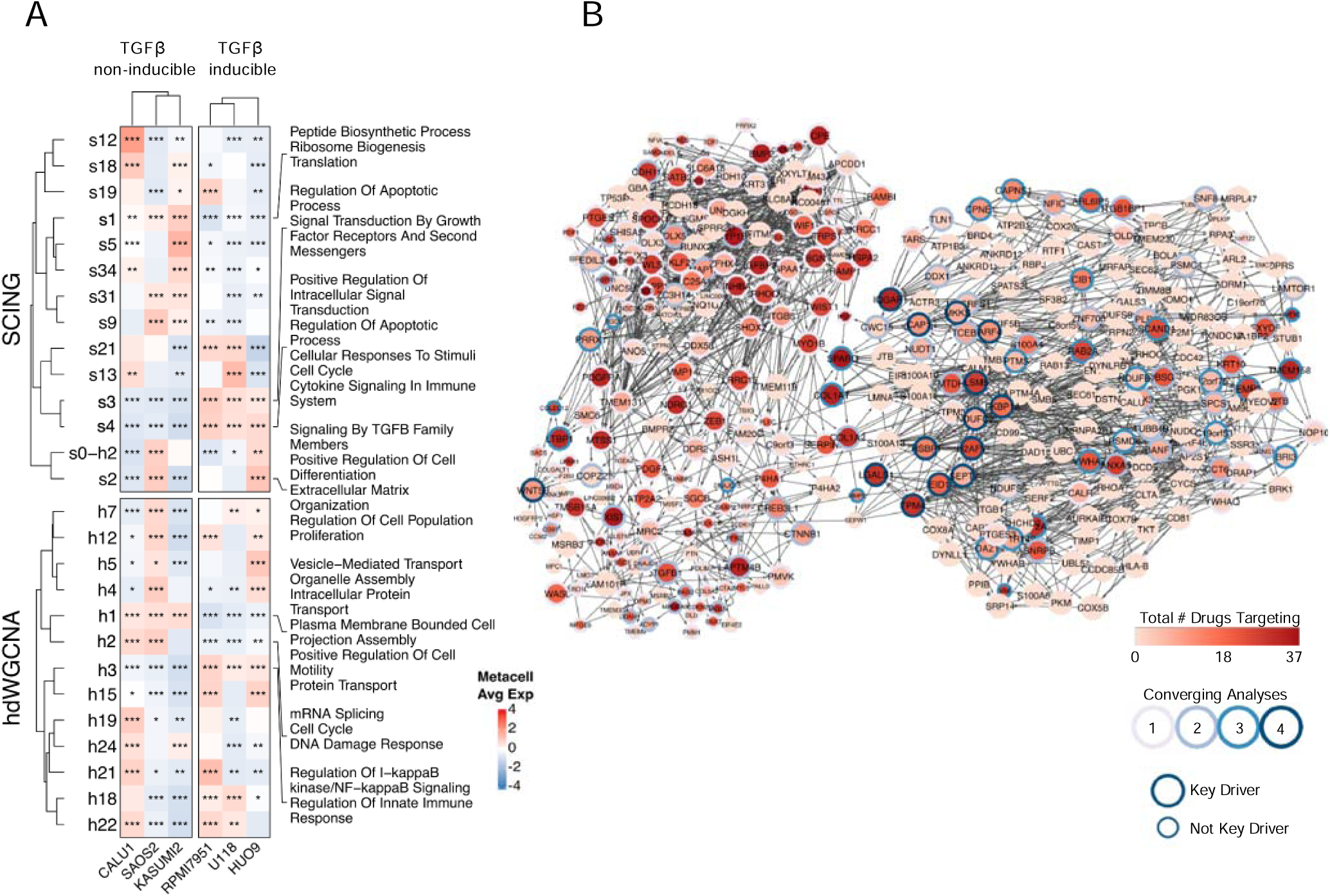
Network Analysis. **(A)** Differential module expression heatmap. Cell lines are grouped by TGFB inducibility. Significance measured by one-vs-opposite group (*<0.05, **<0.01, ***<0.001). **(B)** GRN of top drug targets.

**SUPPLEMENTAL FIGURE 3.**
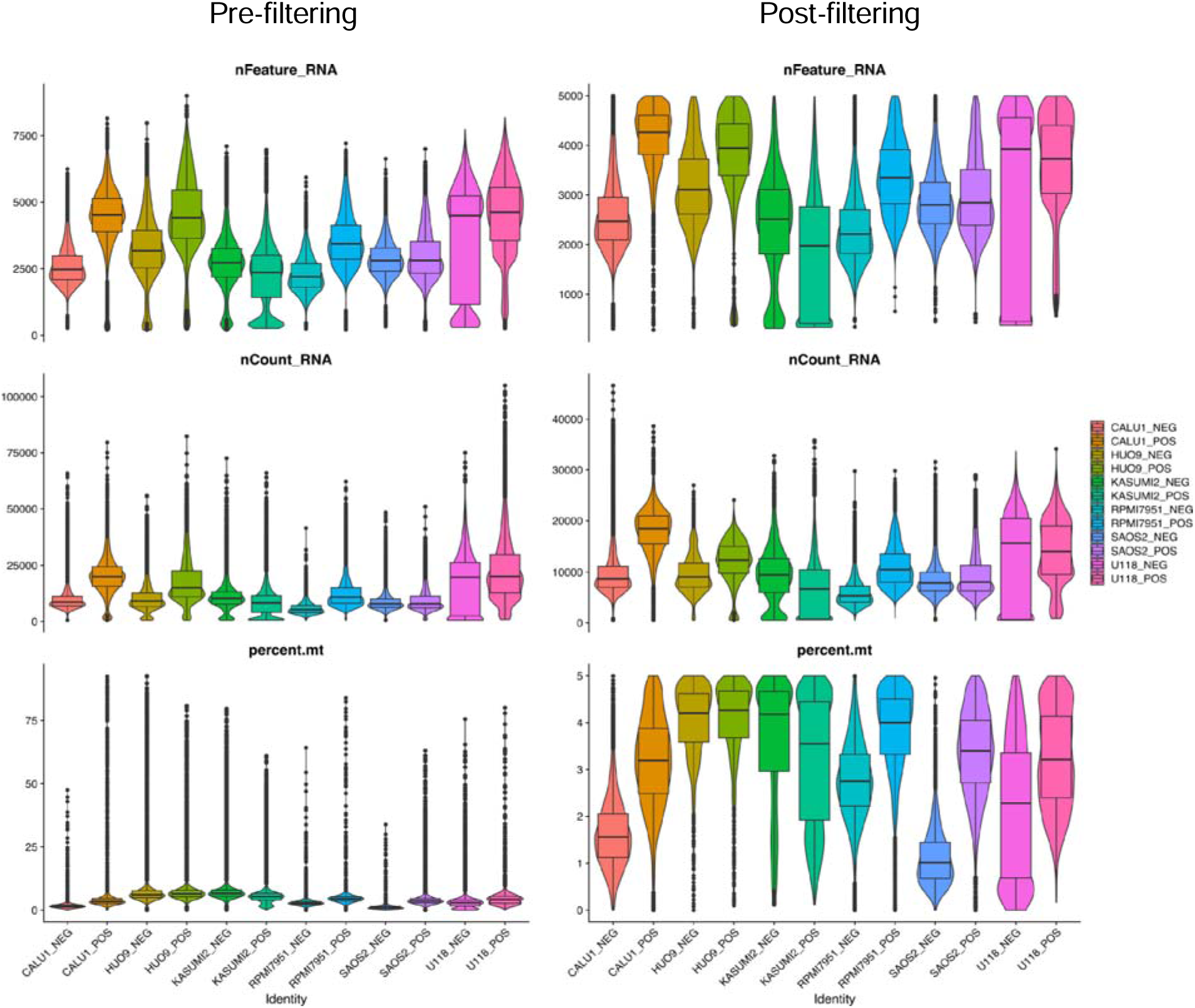
scRNAseq quality control plots pre-filtering (left) and post-filtering (right). Top box plots are the number of genes per cell. Middle box plots are the number of UMI counts per cell. Bottom box plots are the percentage of counts mapping to mitochondrial genes per cell.

**SUPPLEMENTAL FIGURE 4.**
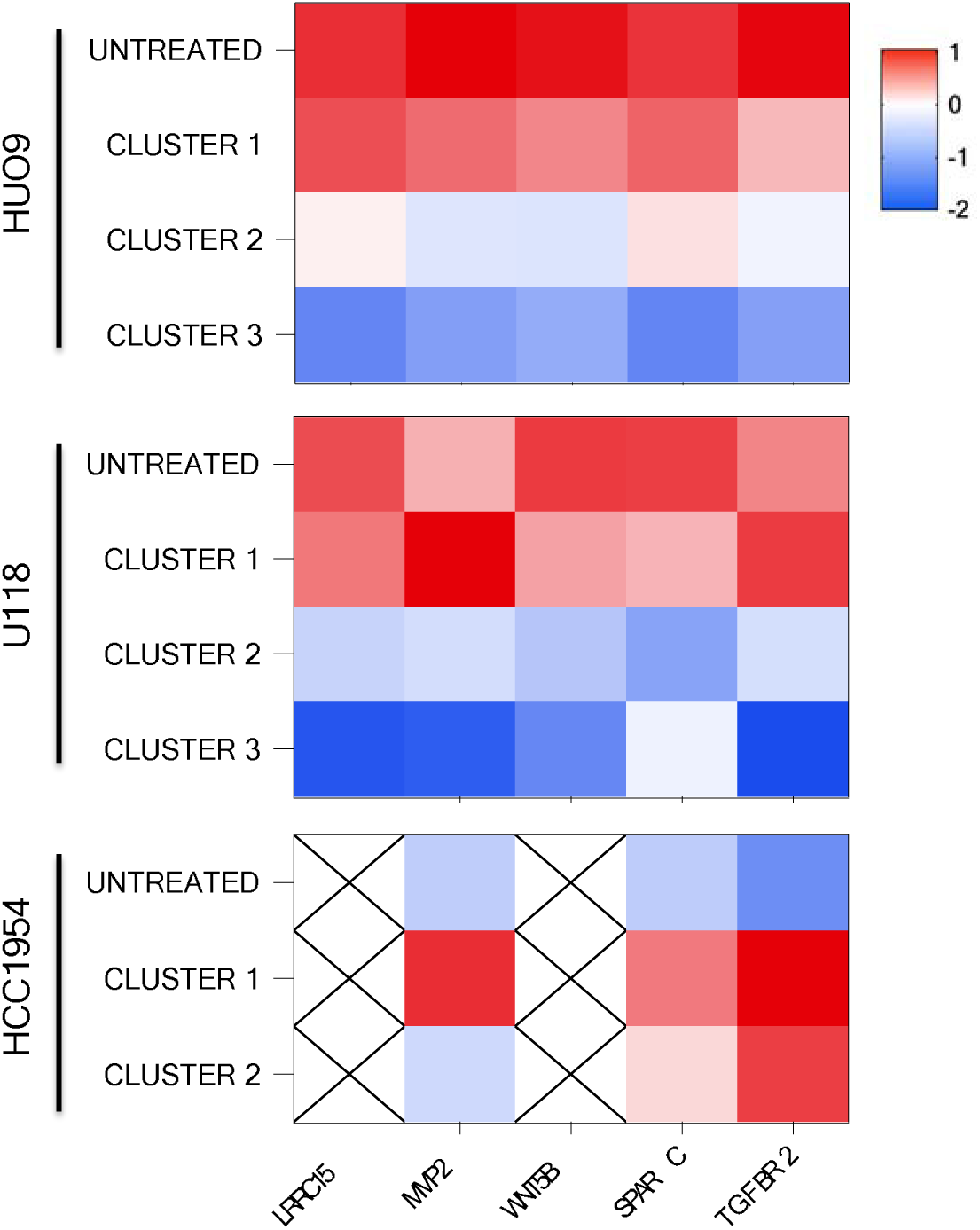
Loss of gene expression in LRRC15 activators after LRRC15-targeted radioimmunotherapy. Z-score normalized transcriptomic expression of *LRRC15*, *MMP2*, *WNT5B*, *SPARC*, and *TGF*β*R2* across [^177^Lu]DUNP19-treated HUO9 (osteosarcoma), U118 (glioblastoma), and HCC1954 (breast carcinoma) tumors (15). High expression (red = high) is observed in untreated tumors, while progressive loss of gene expression is observed in treated tumors (blue = low). Tumors were clustered based on principal component analysis.

**SUPPLEMENTAL FIGURE 5.**
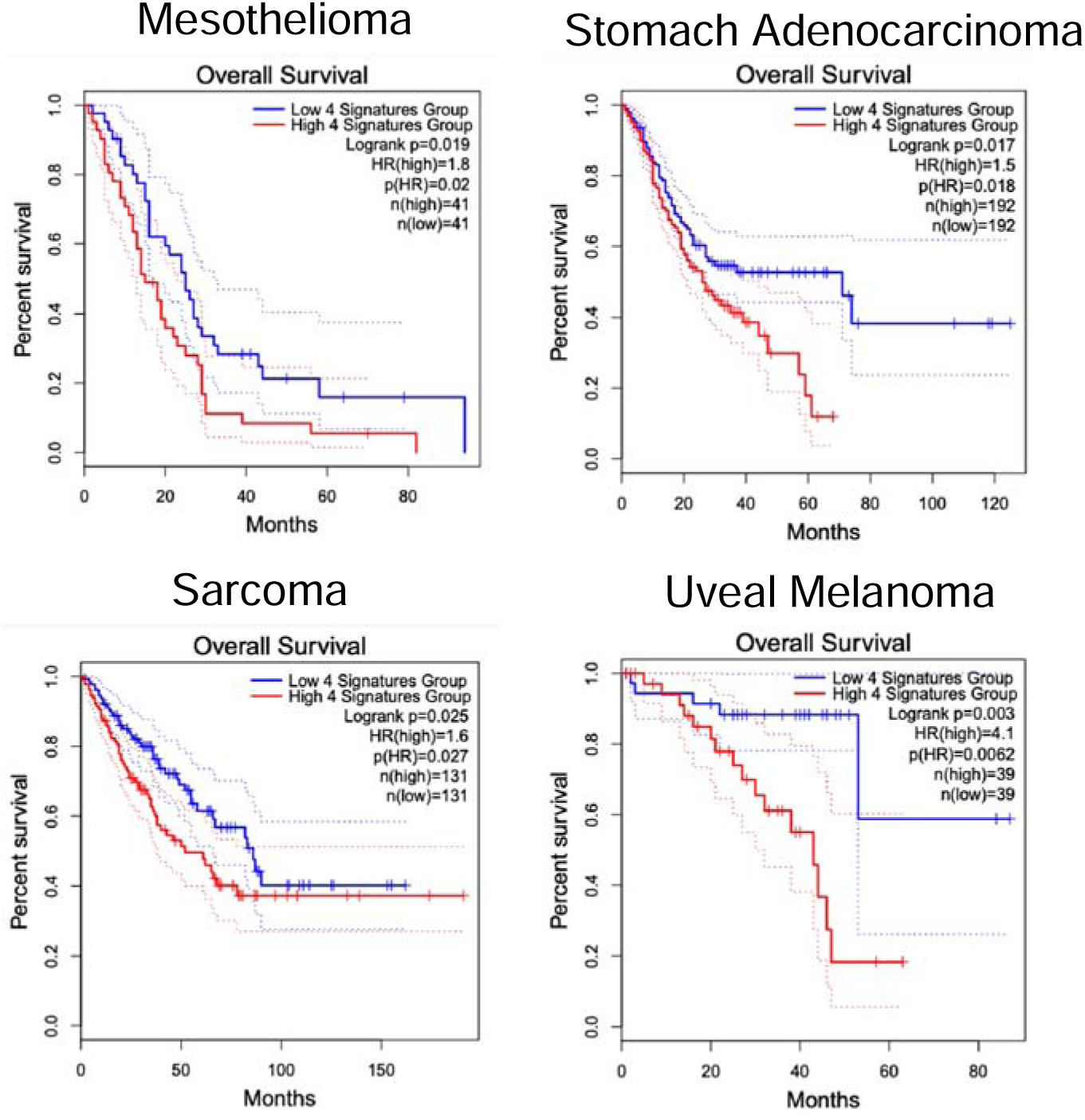
LRRC15 expression activators predict survival in TCGA mesenchymal-derived tumors. TCGA overall survival data comparing patient survival, with high *LRRC15*, *MMP2*, *WNT5B*, *SPARC*, and *TGF*β*R2* gene expression (red) significantly decreasing survival probability across mesenchymal-derived tumor types.

**SUPPLEMENTAL FIGURE 6.**
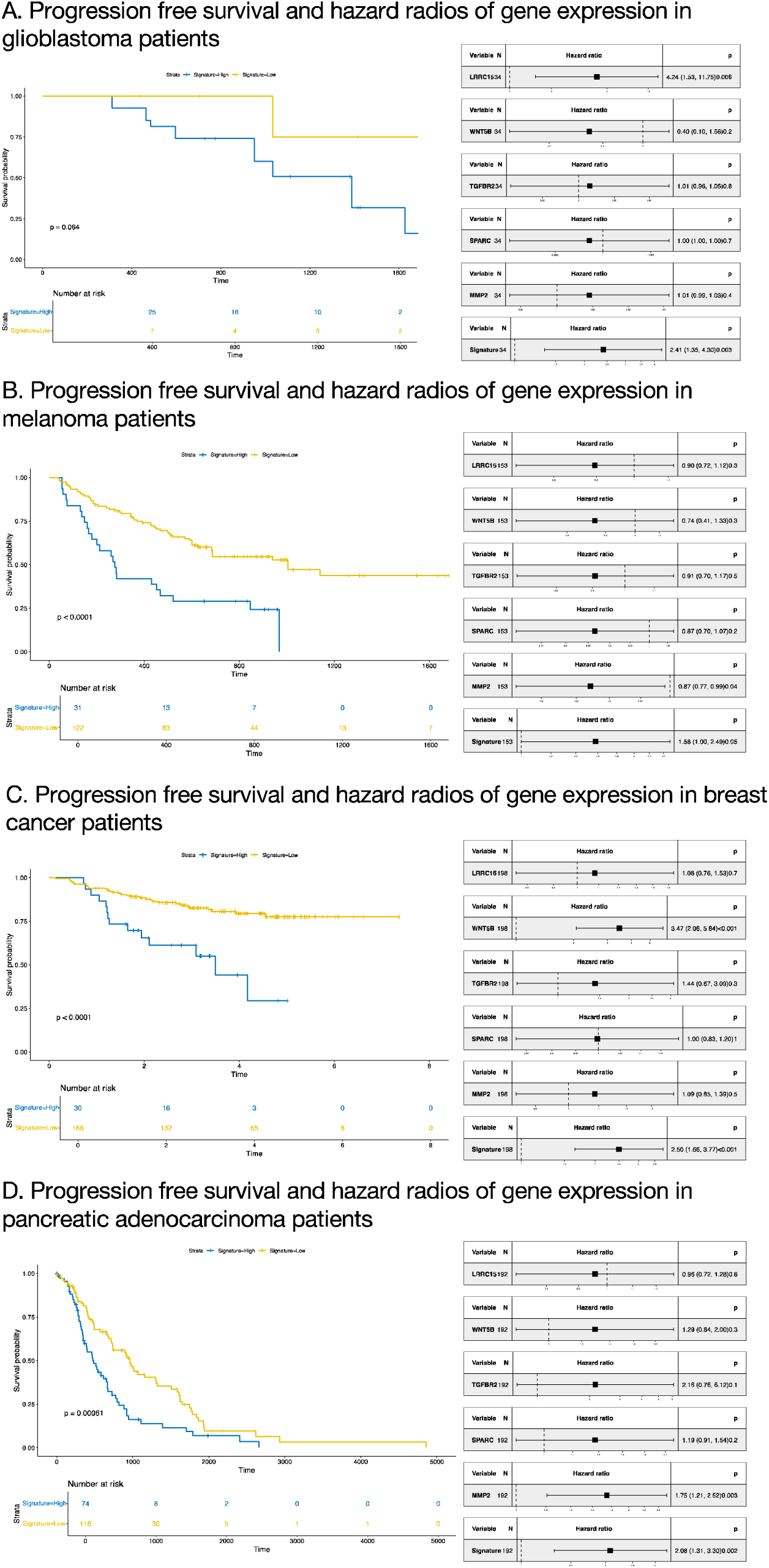
Survival and hazard ratios in tumors based on cell line models. Progression-free survival probability and hazard ratios for individual genes, as well as overall gene expression for *LRRC15*, *MMP2*, *WNT5B*, *SPARC*, and *TGF*β*R2.* For glioblastoma, melanoma, and pancreatic adenocarcinoma patient cohorts, patients were not differentiated by treatment received. Within the breast cancer patient cohort, patients received taxane therapy.

**SUPPLEMENTAL FIGURE 7.**
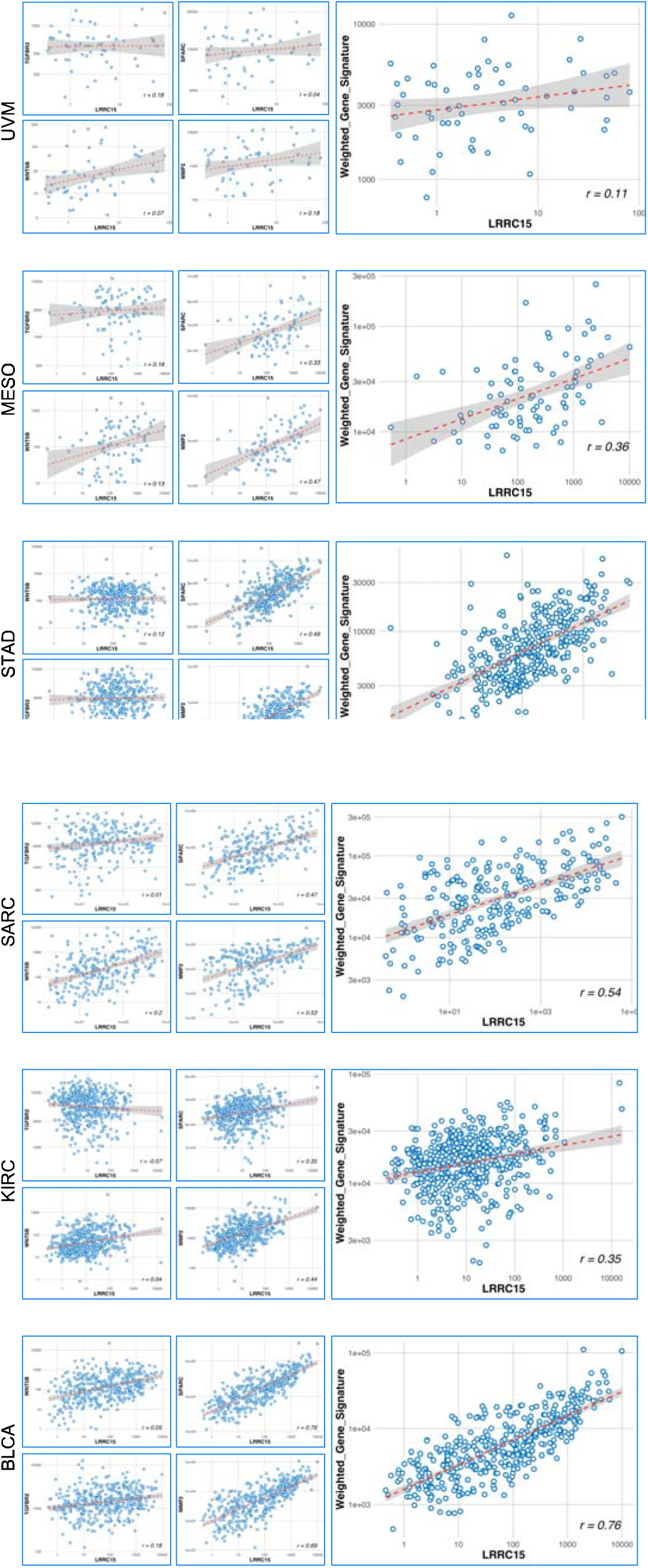
Correlation of LRRC15-associated genes with LRRC15 expression in TCGA tumor cohorts. Correlation plots of uveal melanoma (UVM), mesothelioma (MESO), stomach adenocarcinoma (STAD), sarcoma (SARC), kidney renal cell carcinoma (KIRC) and bladder cancer (BLCA) comparing *MMP2*, *WNT5B*, *SPARC*, and *TGF*β*R2* gene expression to *LRRC15,* either individually (left plots, smaller) or together (right plot, larger). Overall, LRRC15-related genes demonstrate a positive correlation with *LRRC15* expression across tumor types.

**SUPPLEMENTAL FIGURE 8.**
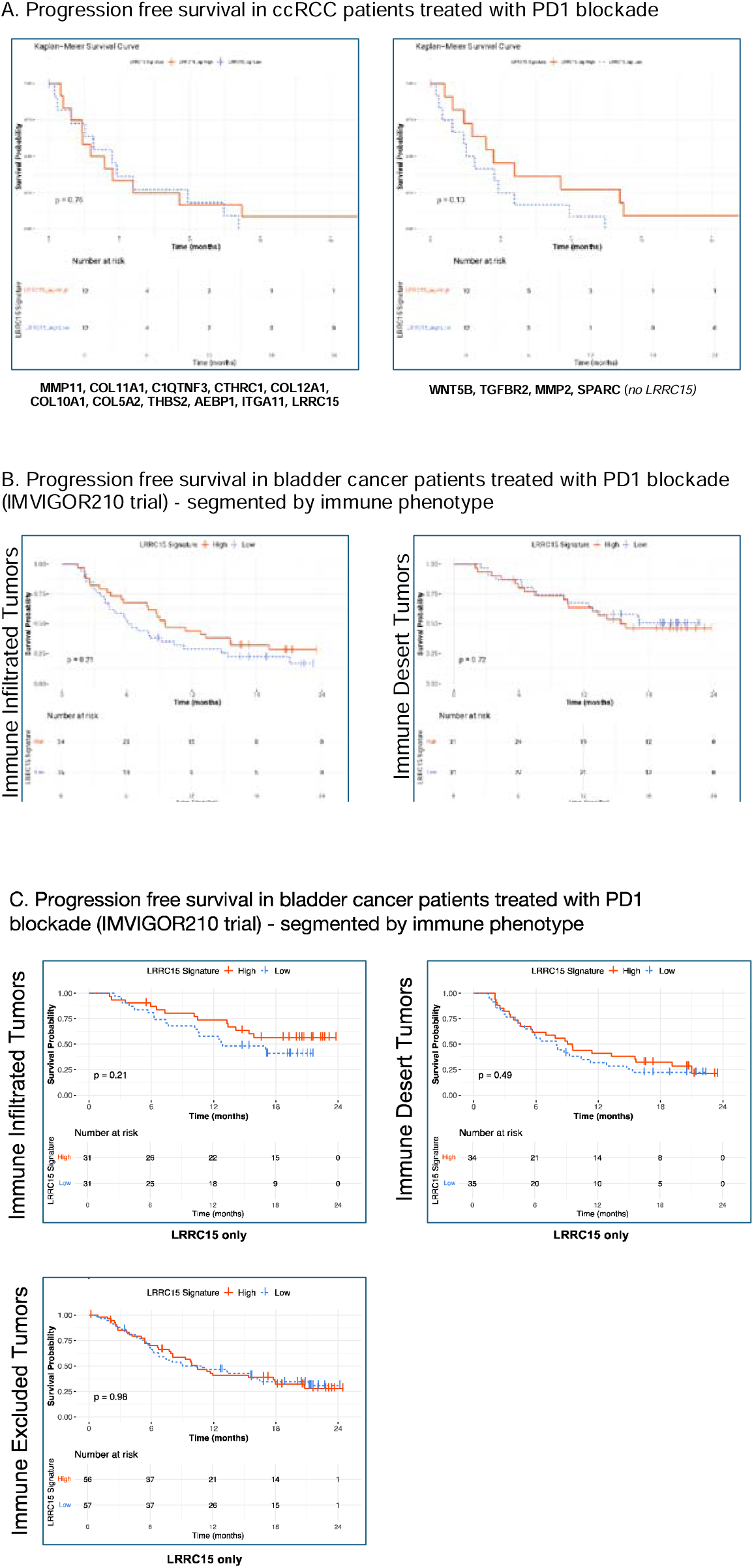
LRRC15 expression activators predict survival in patients with PD1 blockade. **(A)** Progression-free survival in clear cell renal cell carcinoma patients treated with PD1 blockade. Left Kaplan-Meier plot shows no significant predictive ability in signature expression (red = high, blue = low) of a previously identified TGFβ CAF signature that includes LRRC15. Excluding LRRC15 (right plot) decreases predictive power of the LRRC15-related genes *MMP2*, *WNT5B*, *SPARC*, and *TGF*β*R2*.

**SUPPLEMENTAL FIGURE 9.**
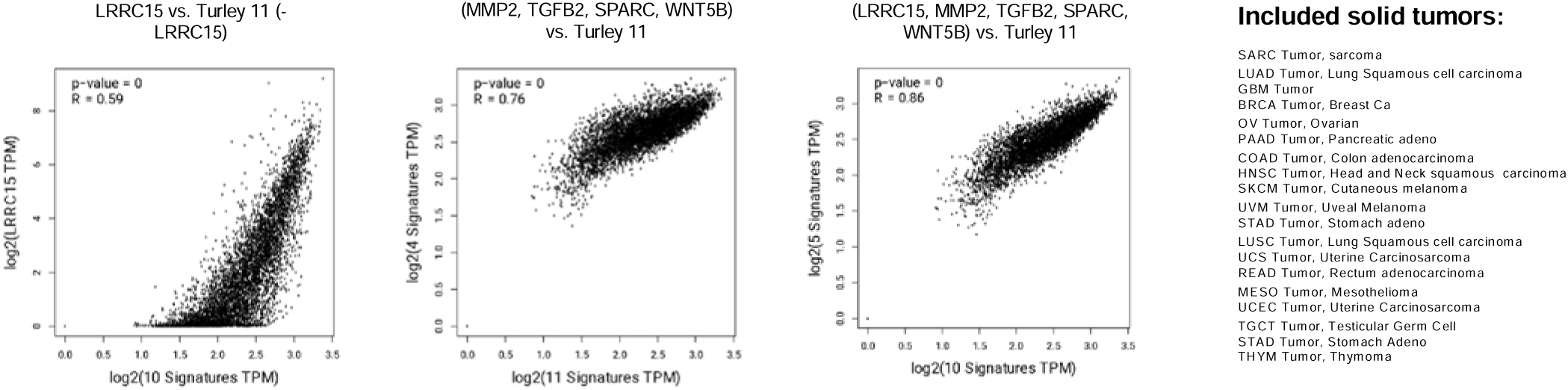
Correlation between previously identified CAF LRRC15 signatures and our LRRC15 gene signatures. (Left) Correlation plots of solid tumor expression of previously identified TGFβ-driven CAF signature by Turley et al. (Turley-11) to LRRC15 expression (left plot). **(Middle)** Plot demonstrates a strong correlation between LRRC15-related genes *MMP2*, *WNT5B*, *SPARC*, and *TGF*β*R2* and the TGFβ-CAF Turley-11 signature. **(Right)** Strong correlation between *MMP2*, *WNT5B*, *SPARC*, and *TGF*β*R2* plus *LRRC15* and the TGFβ-CAF Turley-11 signature.

**SUPPLEMENTAL TABLE 1.** Compound hits from LOPAC1280 screen across three cell lines. Hits are represented as normalized effect on LRRC15 expression.

|  | HUO9 | RPMI7951 | U118 |
| --- | --- | --- | --- |
| (±)-2-Amino-7-phosphonoheptanoic acid | -3.586 |  |  |
| (±)-Quinpirole dihydrochloride | -3.769 | -3.308 | -3.335 |
| 1-Methylimidazole |  | -3.028 | -3.842 |
| 3-aminobenzamide |  | -3.007 | -4.256 |
| AC-93253 iodide |  |  |  |
| AS-252424 | -3.093 |  |  |
| Auranofin | -6.344 | -3.479 | -3.23 |
| Aurothioglucose | -3.53 |  |  |
| Bay 11-7082 |  |  |  |
| Beclomethasone |  |  |  |
| Betamethasone |  |  |  |
| BIX 01294 trihydrochloride hydrate |  |  |  |
| Brefeldin A | -4.692 |  |  |
| Bromoacetyl alprenolol menthane | -4.534 |  |  |
| Budesonide |  |  |  |
| Chloroquine diphosphate |  |  |  |
| Cilnidipine |  |  |  |
| Colchicine | -3.414 |  |  |
| Dihydroouabain |  | -3.85 | -3.016 |
| Ebastine | -5.693 |  |  |
| Emetine dihydrochloride hydrate | -5.796 |  | -4.387 |
| Gossypol | -6.687 |  | -4.296 |
| GW5074 | -4.242 | -3.774 | -3.025 |
| Hydrocortisone |  | -3.204 | -3.34 |
| Hydroxytacrine maleate |  | -3.142 | -3.38 |
| JX401 | -5.488 |  |  |
| K114 |  |  |  |
| Labetalol hydrochloride |  |  |  |
| MDL 105,519 | -3.254 |  |  |
| N-(3,3-Diphenylpropyl)glycinamide |  | -3.309 | -3.326 |
| N-Oleoyldopamine | -3.976 |  |  |
| Niclosamide |  |  |  |
| NSC 95397 | -3.324 |  |  |
| Ouabain | -4.537 |  |  |
| PD-161570 |  |  |  |
| PD-166285 hydrate | -3.965 |  |  |
| PD-184161 |  | -3.425 | -3.244 |
| PD-407824 | -4.41 | -4.001 | -3.015 |
| PD173952 | -3.436 |  |  |
| Phorbol 12-myristate 13-acetate |  | -3.114 | -3.569 |
| Pifithrin-mu |  | -3.376 | -3.311 |
| Podophyllotoxin | -3.571 |  |  |
| RepSox | -5.36 | -3.603 | -3.035 |
| Rottlerin |  | -3.262 | -3.34 |
| Sanguinarine chloride |  | -3.491 | -3.171 |
| SB-525334 | -5.045 | -3.803 | -3.022 |
| Stattic | -5.231 |  |  |
| TBB |  | -3.577 | -3.038 |
| Terfenadine |  |  |  |
| **Thapsigargin** | -4.334 | -3.413 | -3.257 |
| **Topotecan hydrochloride hydrate** | -5.435 |  |  |
| **Triamcinolone** |  |  |  |
| **TTNPB** |  | -3.399 | -3.278 |
| **Tyrphostin AG 1478** |  | -3.179 | -3.354 |
| **Tyrphostin AG 879** |  | -3.494 | -3.12 |
| **Total Compound Hits** | **26** | **22** | **24** |

## REFERENCES

1. Bierie, B. & Moses, H. L. Tumour microenvironment: TGFbeta: the molecular Jekyll and Hyde of cancer. Nat Rev Cancer 6, 506–520 (2006).

2. Biswas, T., Gu, X., Yang, J., Ellies, L. G. & Sun, L.-Z. Attenuation of TGF-β signaling supports tumor progression of a mesenchymal-like mammary tumor cell line in a syngeneic murine model. Cancer Lett 346, 129– 138 (2014).

3. Massagué, J. TGFβ signalling in context. Nat Rev Mol Cell Biol 13, 616–630 (2012).

4. Tolcher, A. W. et al. A phase 1 study of anti-TGFβ receptor type-II monoclonal antibody LY3022859 in patients with advanced solid tumors. Cancer Chemother Pharmacol 79, 673–680 (2017).

5. Hezel, A. F. et al. TGF-β and αvβ6 integrin act in a common pathway to suppress pancreatic cancer progression. Cancer Res 72, 4840–4845 (2012).

6. Brandes, A. A. et al. A Phase II randomized study of galunisertib monotherapy or galunisertib plus lomustine compared with lomustine monotherapy in patients with recurrent glioblastoma. Neuro Oncol 18, 1146–1156 (2016).

7. Yang, Y. et al. The Outcome of TGFβ Antagonism in Metastatic Breast Cancer Models In Vivo Reflects a Complex Balance between Tumor-Suppressive and Proprogression Activities of TGFβ. Clin Cancer Res 26, 643– 656 (2020).

8. Connolly, E. C. et al. Outgrowth of drug-resistant carcinomas expressing markers of tumor aggression after long-term TβRI/II kinase inhibition with LY2109761. Cancer Res 71, 2339–2349 (2011).

9. Derynck, R., Turley, S. J. & Akhurst, R. J. TGFβ biology in cancer progression and immunotherapy. Nat Rev Clin Oncol 18, 9–34 (2021).

10. Cui, J. et al. Expression and clinical implications of leucine-rich repeat containing 15 (LRRC15) in osteosarcoma. J Orthop Res 38, 2362–2372 (2020).

11. Ray, U. et al. Exploiting LRRC15 as a Novel Therapeutic Target in Cancer. Cancer Res 82, 1675–1681 (2022).

12. Ray, U. et al. Targeting LRRC15 Inhibits Metastatic Dissemination of Ovarian Cancer. Cancer Res 82, 1038– 1054 (2022).

13. Krishnamurty, A. T. et al. LRRC15(+) myofibroblasts dictate the stromal setpoint to suppress tumour immunity. Nature 611, 148–154 (2022).

14. Dominguez, C. X. et al. Single-Cell RNA Sequencing Reveals Stromal Evolution into LRRC15(+) Myofibroblasts as a Determinant of Patient Response to Cancer Immunotherapy. Cancer Discov 10, 232–253 (2020).

15. Storey, C. M. et al. Development of a LRRC15-Targeted Radio-Immunotheranostic Approach to Deplete Pro-tumorigenic Mechanisms and Immunotherapy Resistance. Preprint at https://www.biorxiv.org/content/10.1101/2024.01.30.577289v1 (2024).

16. Purcell, J. W. et al. LRRC15 Is a Novel Mesenchymal Protein and Stromal Target for Antibody-Drug Conjugates. Cancer Res 78, 4059–4072 (2018).

17. Wakefield, L. M. et al. Transforming growth factor-beta1 circulates in normal human plasma and is unchanged in advanced metastatic breast cancer. Clin Cancer Res 1, 129–136 (1995).

18. Shilts, J. et al. LRRC15 mediates an accessory interaction with the SARS-CoV-2 spike protein. PLoS Biol 21, e3001959 (2023).

19. Morabito, S., Reese, F., Rahimzadeh, N., Miyoshi, E. & Swarup, V. hdWGCNA identifies co-expression networks in high-dimensional transcriptomics data. Cell Rep Methods 3, 100498 (2023).

20. Barata, J. T., Durum, S. K. & Seddon, B. Flip the coin: IL-7 and IL-7R in health and disease. Nat Immunol 20, 1584–1593 (2019).

21. Littman, R., Cheng, M., Wang, N., Peng, C. & Yang, X. SCING: Inference of robust, interpretable gene regulatory networks from single cell and spatial transcriptomics. iScience 26, 107124 (2023).

22. Traag, V. A., Waltman, L. & van Eck, N. J. From Louvain to Leiden: guaranteeing well-connected communities. Sci Rep 9, 5233 (2019).

23. Subramanian, A. et al. A Next Generation Connectivity Map: L1000 Platform and the First 1,000,000 Profiles. Cell 171, 1437–1452.e17 (2017).

24. Ghosh, A. K. & Varga, J. The transcriptional coactivator and acetyltransferase p300 in fibroblast biology and fibrosis. J Cell Physiol 213, 663–671 (2007).

25. Cerami, E. et al. The cBio cancer genomics portal: an open platform for exploring multidimensional cancer genomics data. Cancer Discov 2, 401–404 (2012).

26. A Study of Atezolizumab in Participants With Locally Advanced or Metastatic Urothelial Bladder Cancer (Cohort 2). Clinicaltrials.gov identifier: NCT02108652

27. Cook, D. et al. Lessons learned from the fate of AstraZeneca’s drug pipeline: a five-dimensional framework. Nat Rev Drug Discov 13, 419–431 (2014).

28. Emmerich, C. H. et al. Improving target assessment in biomedical research: the GOT-IT recommendations. Nat Rev Drug Discov 20, 64–81 (2021).

29. Tsellou, E. & Kiaris, H. Fibroblast independency in tumors: implications in cancer therapy. Future Oncol 4, 427– 432 (2008).

30. Hill, R., Song, Y., Cardiff, R. D. & Van Dyke, T. Selective evolution of stromal mesenchyme with p53 loss in response to epithelial tumorigenesis. Cell 123, 1001–1011 (2005).

31. Littlepage, L. E., Egeblad, M. & Werb, Z. Coevolution of cancer and stromal cellular responses. Cancer Cell 7, 499–500 (2005).

32. Olive, K. P. et al. Inhibition of Hedgehog signaling enhances delivery of chemotherapy in a mouse model of pancreatic cancer. Science 324, 1457–1461 (2009).

33. Carvalheiro, T. et al. Extracellular SPARC cooperates with TGF-β signalling to induce pro-fibrotic activation of systemic sclerosis patient dermal fibroblasts. Rheumatology (Oxford) 59, 2258–2263 (2020).

34. Fan, J. et al. SPARC knockdown attenuated TGF-β1-induced fibrotic effects through Smad2/3 pathways in human pterygium fibroblasts. Arch Biochem Biophys 713, 109049 (2021).

35. Chew, G.-L. et al. Short H2A histone variants are expressed in cancer. Nat Commun 12, 490 (2021).

36. Lamaa, A. et al. Integrated analysis of H2A.Z isoforms function reveals a complex interplay in gene regulation. Elife 9, (2020).

37. Leung, J. W. C., Emery, L. E. & Miller, K. M. CRISPR/Cas9 Gene Editing of Human Histone H2A Variant H2AX and MacroH2A. Methods Mol Biol 1832, 255–269 (2018).

38. Boettcher, M. & McManus, M. T. Choosing the Right Tool for the Job: RNAi, TALEN, or CRISPR. Mol Cell 58, 575–585 (2015).

39. Pembrolizumab and Dasatinib, Imatinib Mesylate, or Nilotinib in Treating Patients With Chronic Myeloid Leukemia and Persistently Detectable Minimal Residual Disease. Clinicaltrials.gov identifier: NCT03516279

40. Xuan, J., Yu, Y., Qing, T., Guo, L. & Shi, L. Next-generation sequencing in the clinic: promises and challenges. Cancer Lett 340, 284–295 (2013).

41. Flaherty, K. T. et al. The Molecular Analysis for Therapy Choice (NCI-MATCH) Trial: Lessons for Genomic Trial Design. J Natl Cancer Inst 112, 1021–1029 (2020).

42. Flaherty, K. T. et al. Molecular Landscape and Actionable Alterations in a Genomically Guided Cancer Clinical Trial: National Cancer Institute Molecular Analysis for Therapy Choice (NCI-MATCH). J Clin Oncol 38, 3883– 3894 (2020).

43. Meric-Bernstam, F. et al. National Cancer Institute Combination Therapy Platform Trial with Molecular Analysis for Therapy Choice (ComboMATCH). Clin Cancer Res 29, 1412–1422 (2023).

44. Hao, Y. et al. Integrated analysis of multimodal single-cell data. Cell 184, 3573–3587.e29 (2021).

45. Satija, R., Farrell, J. A., Gennert, D., Schier, A. F. & Regev, A. Spatial reconstruction of single-cell gene expression data. Nat Biotechnol 33, 495–502 (2015).

46. Shu, L. et al. Mergeomics: multidimensional data integration to identify pathogenic perturbations to biological systems. BMC Genomics 17, 874 (2016).

47. Kuleshov MV, Jones MR, Rouillard AD, et al. Enrichr: a comprehensive gene set enrichment analysis web server 2016 update. Nucleic Acids Res. 2016;44(W1):W90–W97.

48. Shannon P, Markiel A, Ozier O, et al. Cytoscape: a software environment for integrated models of biomolecular interaction networks. Genome Res. 2003;13(11):2498–2504.

